# Plasmids shape the diverse accessory resistomes of *Escherichia coli* ST131

**DOI:** 10.1101/2020.05.07.081380

**Authors:** Arun Gonzales Decano, Nghia Tran, Hawriya Al-Foori, Buthaina Al-Awadi, Leigh Campbell, Kevin Ellison, Louisse Paola Mirabueno, Maddy Nelson, Shane Power, Genevieve Smith, Cian Smyth, Zoe Vance, Caitriona Woods, Alexander Rahm, Tim Downing

## Abstract

The human gut microbiome includes beneficial, commensal and pathogenic bacteria that possess antimicrobial resistance (AMR) genes that exchange these predominantly through conjugative plasmids. *Escherichia coli* is a significant component of the gastrointestinal microbiome and is typically non-pathogenic in this niche. In contrast, extra-intestinal pathogenic *E. coli* (ExPEC) including ST131 may occupy other environments like the urinary tract or bloodstream where they express genes enabling AMR and host cell adhesion like type 1 fimbriae. The extent to which commensal *E. coli* and uropathogenic ExPEC ST131 share AMR genes remains understudied at a genomic level, and here we examined this using a preterm infant resistome. Here, individual ST131 had small differences in AMR gene content relative to a larger shared resistome. Comparisons with a range of plasmids common in ST131 showed that AMR gene composition was driven by conjugation, recombination and mobile genetic elements. Plasmid pEK499 had extended regions in most ST131 Clade C isolates, and it had evidence of a co-evolutionary signal based on protein-level interactions with chromosomal gene products, as did pEK204 that had a type IV fimbrial *pil* operon. ST131 possessed extensive diversity of selective type 1, type IV, P and F17-like fimbriae genes that was highest in subclade C2. The structure and composition of AMR genes, plasmids and fimbriae vary widely in ST131 Clade C and this may mediate pathogenicity and infection outcomes.

**Data Summary:** The following files are available on the FigShare project “Plasmids_ST131_resistome_2020” :

1. The set of 794 AMR genes derived from [74] are available (with their protein sequence translation) at FigShare at doi: dx.doi.org/10.6084/m9.figshare.11961402.
2. The AMR gene profiles per sample determined by their BLAST sequence similarity results against CARD are available at FigShare at doi: dx.doi.org/10.6084/m9.figshare.11961612. This dataset includes the PlasmidFinder results. It also includes other AMR database comparisons (ARG-ANNOT, ResFinder, MegaRes, VFDB and VirulenceFinder).
3. The BLAST sequence similarity results for the *fim, pil, pap* and *ucl* operons’ genes versus 4,071 *E. coli* ST131 assemblies from Decano & Downing (2019) are available at FigShare at doi: dx.doi.org/10.6084/m9.figshare.11961711.
4. The genome sequences and annotation files for reference genomes NCTC13441, EC958 and SE15, along with the assembled contigs for 83972 and 3_2_53FAA are available at FigShare at doi: dx.doi.org/10.6084/m9.figshare.11961813.
5. The 4,071 *E. coli* ST131 genome assemblies from Decano & Downing (2019) are available at FigShare at doi: 10.6084/m9.figshare.11962278 (the first 1,680 assemblies) and at doi: dx.doi.org/10.6084/m9.figshare.11962557 (the second 2,391 assemblies).

## Introduction

Extra-intestinal pathogenic *E. coli* (ExPEC) cause extensive infections outside the gut, from which they can originate. ExPEC have a wide range of virulence factor (VF) [1-4] and antimicrobial resistance (AMR) genes [5], especially sequence type (ST) 131 from serotypes O25b:H4 and O16:H5 in phylogroup B2 [6]. ST131 causes a substantial fraction of ExPEC and extended-spectrum beta-lactamase (ESBL)-producing cases [7]. ST131’s acquisition of key AMR genes encoded on plasmids [4,8] has coincided with its adaptation to new environments [9,10].

ExPEC adherence factors allowing colonisation of different host niches are mainly produced by the Chaperone-Usher secretion pathway, including type 1, P, F1C/S and AFA fimbriae [11-14]. Within ExPEC ST131, fluoroquinolone-resistant Clade C is the main cause of human infection globally [15-16]. Its D-mannose-specific type 1 fimbriae encoded by the *fim* operon binds the host mucosal epithelium to cause urinary tract and kidney infection [18-21]. The genetic diversity of fimbrial operons beyond *fim* in Clade C has not yet been examined in large collections of isolates.

ExPEC infection can be associated with a changed microbiome composition [22]. *E. coli* is the most common initial coloniser of infant intestines [23], where most are commensal [24] and some protect against pathogen invasion [25]. Nonetheless, AMR is prevalent in neonatal units [26] and within-host gene exchange between commensal and pathogenic bacteria may occur [27]. ST131 spreads between mother-infant pairs [28] likely via an oro-faecal transmission route [29], and such colonisation of infants can last for long periods [30]. Consequently, AMR gene screening can inform on treatment strategies [31] and the effect of antibiotics on microbiomes [32].

The ST131 resistome (the set of AMR genes) includes ESBL genes [33-34] allowing third-generation cephalosporin-resistance [35] and are associated with three main cefotaximase (CTX-M) resistance alleles: *bla*_CTX-M-14/15/27_ [36]. Like most AMR genes, these are sandwiched by mobile genetic elements (MGEs) on plasmids and thus can be gained by horizontal gene transfer (HGT) [37-39] or lost if not beneficial [40-41]. Of nine common bacterial pathogens, *E. coli* has the most MGEs, including phage-associated ones and transpose (*tnpA*) genes [42]. Diverse pathogenic bacteria share MGEs [43-45] and HGT may worsen nosocomial outbreaks [46-48]. Plasmids common to ST131 often have IS*Ecp1*, IS*903D* and IS*26* elements [49] encoding *tnpA* flanked by short inverted repeats, typically with ESBL genes. Such ESBL genes can be chromosomally transferred and may form part of the core resistome [50].

Fluctuating antibiotic exposure, host type, anatomical niche and immunity mean that resistomes vary [51]. Conjugation, recombination and MGEs -help shape ST131 resistome dynamics [1,10,52-55]. ST131’s plasmids typically are from incompatibility (Inc) groups F, I1, N and A/C [56-58]. Some plasmids in ST131 encode genes for post-segregation killing and stable inheritance to ensure their propagation, but these genes can be lost or may recombine with other plasmids [58-60]. As a result of this mixing and their extensive array of MGEs, plasmids may rearrange extensively even within a clonal radiation [61-62]. Plasmids may also impair cell reproduction due to the energetic cost of their replication and maintenance, so conjugation and recombination could allow gene dosage optimisation and gene expression coordination.

There is extensive evidence of AMR gene conjugation in the gut between species including *E. coli* [63-66], and also between its STs, such as: transfer of a *bla*_CTX-M-1_-positive Incl1 plasmid among ST1640, ST2144 and ST6331 in a single patient’s gut [67]; a *bla*_OXA-48_ gene on a *K. pneumoniae* IncL/M-type plasmid from ST14 to ST666 [68] and from ST648 to ST866 [69]; a 113.6 kb IncF plasmid with a mercury detoxification operon *bla*_KPC-2_, *bla*_OXA-9_ and *bla*_TEM-1A_ genes from ST69 to ST131 [70]; and transfer of a range of sulphonamide-(*sul2*) and ampicillin-resistance genes (*bla*_TEM-1b_) on a pNK29 plasmid within *E. coli* subtypes [71-72]. Moreover, the frequency of conjugation was ten times higher and more stable in *bla*_CTX-M-15_-producing ST131 than in *K. pneumoniae* [73].

This study focused on uropathogenic ST131 Clade C (*fimH30*) because of its high plasmid and AMR gene diversity [50,61] where the mobile resistome and how plasmids mediate AMR gene composition needs deeper scrutiny. Here, we found a large shared core preterm infant resistome and a smaller accessory one that varied between closely related isolates. This was caused by plasmid turnover and rearrangements, affecting fimbrial genes too. We hypothesise that certain plasmids are more compatible with ST131 genomes, and applied topological data analysis (TDA) to understand plasmid-chromosome co-evolution.

## Results

### A large core preterm infant resistome in *E. coli*

We examined the preterm infant resistome and plasmid composition of seven *E. coli* ST131 genomes in the context of related reference genomes and non-pathogenic isolates. This resistome of 794 genes was assembled from 2,004 contigs originally derived from DNA sequencing of 21 preterm infant faecal samples experimentally tested *in vitro* for resistance to 16 antibiotics (see Data Access, Table S1) [74]. This resistome provided a culture-unbiased perspective on antibiotic resistance genes [74]. We found that seven ST131 Clade C genome assemblies from adult urinary tract infections (UTIs) together with subclade C2 reference genomes EC958 and NCTC13441 had 262 to 291 unique contigs in this resistome (Table 1). Commensal ST131 Clade A reference SE15 (O150:H5, *fimH41*) had 251 AMR contigs, and intestinal *E. coli* Human Microbiome Project (HMP) assembly 3_2_53FAA had 269. In contrast, well-studied asymptomatic urinary tract HMP sample 83972 from ST73 [75] had 213 unique contigs, suggesting a subset of AMR genes may be associated with symptomatic ST131 UTIs.

**Table 1.** *E. coli* genome assembly collection and metadata. The table shows the sample ID, SRA accession ID, ST131 Clade, *bla*_CTX-M_ allele(s), country and year of isolation, and numbers of unique AMR contigs in the preterm infant resistome. All seven ST131 assemblies, EC958 and NCTC13441 came from pathogenic UTIs. SE15 was a faecal commensal isolate. EC958 had 18 AMR contigs on plasmid pEC958A (pEC958B had none). NCTC13441 had 27 AMR contigs on its plasmid. The seven ST131 and two HMP samples were assembled from Illumina reads, and the three references were assembled from PacBio reads.

| Sample ID | SRA ID | ST131<br>clade or ST | *bla*<sub>CTX-M</sub><br>allele(s) | Country, year | Unique<br>AMR contigs |
| --- | --- | --- | --- | --- | --- |
| 8289_1#91 | ERR191724 | C1 | 14 | Ireland, 2007 | 268 |
| 8289_1#35 | ERR191668 | C1 | 14 | Ireland, 2008 | 270 |
| 8289_1#3 | ERR191636 | C2 | 15 | Ireland, 2005 | 274 |
| 8289_1#34 | ERR191667 | C2 | 15 | Ireland, 2010 | 274 |
| 8289_1#24 | ERR191657 | C2 | 14 & 15 | Ireland, 2009 | 283 |
| 8289_1#60 | ERR191693 | C2 | 15 | Ireland, 2010 | 262 |
| 8289_1#27 | ERR191660 | C2 | 15 | Ireland, 2010 | 262 |
| SE15 | AP009378 | A | none | Japan, 2010 | 251 |
| NCTC13441 | ERS530440 | C2 | 15 | UK, 2003 | 280 |
| EC958 | HG941718 | C2 | 15 | UK, 2005 | 291 |
| 3_2_53FAA | - | ST95 | none | Canada, 2007 | 269 |
| 83972 | - | ST73 | none | Sweden, 1978 | 213 |

Comparing the AMR contigs’ gene product functions identified a core resistome of 210 AMR contigs shared by all (Figure 1), highlighting that these may be essential for non-pathogenic isolates. This core resistome differed from the accessory resistome of 109 contigs, of which 50 in ST131 Clade C alone corresponded to 18 unique AMR genes (Figure 1). These accessory AMR genes were identified with ARIBA v2.11.1 [76] and the Comprehensive Antibiotic Resistance Database (CARD) [77], where matches were verified by read mapping to the reference resistome using GROOT [78].

**Figure 1.**
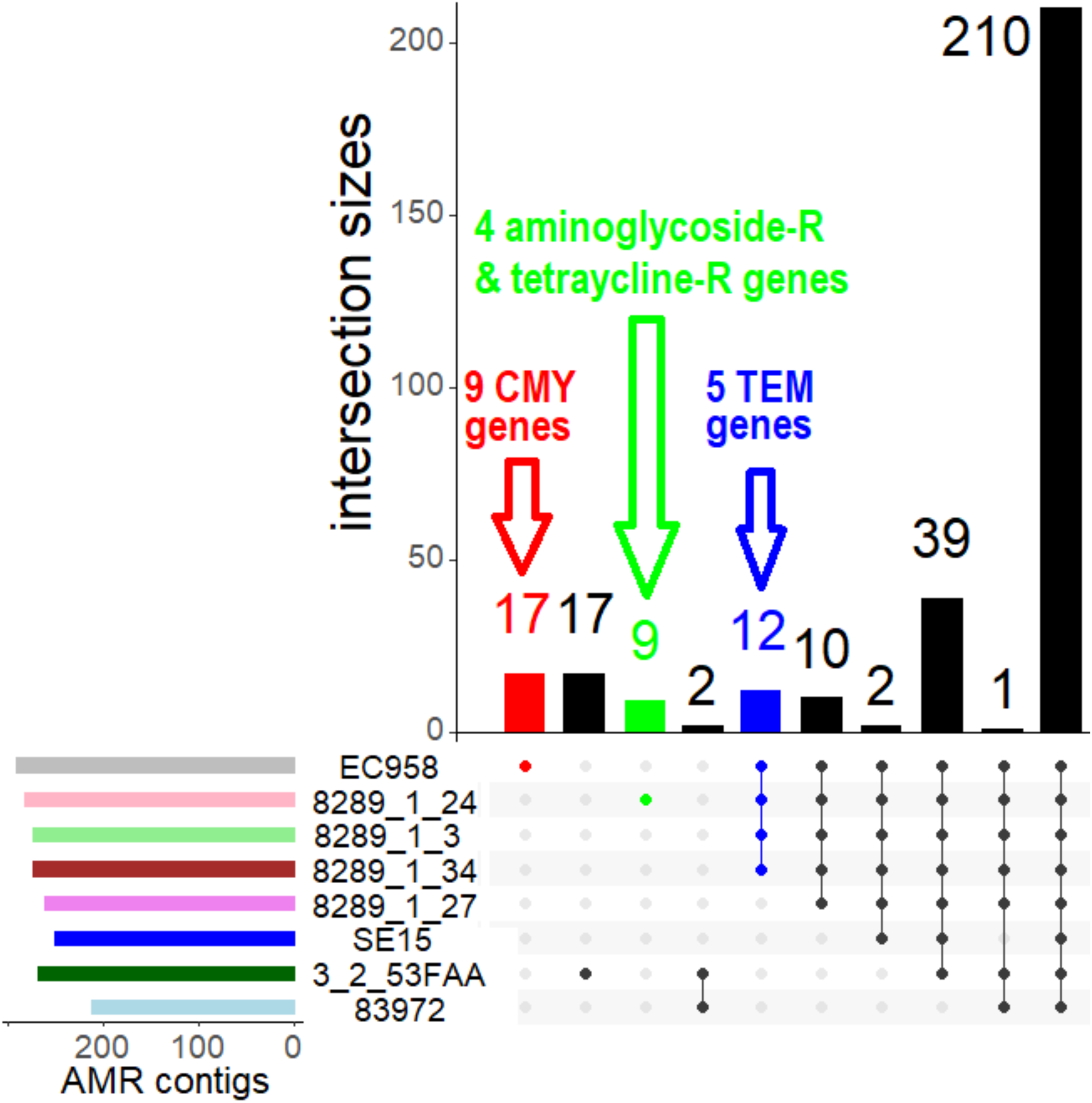
The overlap of preterm infant AMR contigs across four ST131 subclade C2 assemblies (8289_1#3, 8289_1#24, 8289_1#27, 8289_1#34), two ST131 reference genomes (EC958 and SE15) and two HMP assemblies (83972 and 3_2_53FAA). Top: The intersection sizes (y-axis) and the numbers of AMR contigs per set showed that most (210) AMR contigs were shared across all isolates. The non-unique AMR contigs indicated smaller numbers of unique AMR genes, so EC958’s 17 AMR contigs (red) corresponded to nine unique *bla*_CMY_ genes (Table S2); and 8289_1#24’s nine contigs (green) represented four aminoglycoside and tetracycline resistance genes (Table S3). All Clade C bar 8289_1#27 had 12 contigs (blue) denoting to five *bla*_TEM_ genes (Table S4). Bottom: The numbers of AMR contigs per sample with the corresponding sets (coloured circles).

### A diverse accessory preterm infant resistome among closely related ST131 isolates

*Bla*_CMY_ encoding ESBL AmpC is typically associated with cephalosporin-resistance [79] and here nine variants (alleles CMY-37, 51, 66, 67, 85, 98, 101, 105) resolved from 17 AMR contig matches were unique to EC958 (Figure 1). All Subclade C2 isolates bar 8289_1#27 had five *bla*_TEM_ alleles (TEM-57, 104, 116, 215, 220) encoding an Ambler class A β-lactamase. The sole *bla*_CTX-M-14_- and *bla*_CTX-M-15_-positive isolate (8289_1#24) had unique four genes linked to aminoglycoside and tetracycline resistance (Table S2), a combination that may neutralise the effect of aminoglycoside susceptibility associated with tetracycline efflux pump TetA expression [80-81].

Additional comparisons using the NCTC13441 genome showed that it had five *bla*_OXY_ alleles (OXY-1-1, 1-2, 2-3, 6-2, 6-4) encoding an Ambler class A ESBL (Text S1). Ten AMR contigs in all C2 corresponded to a single helix-turn-helix-like transcriptional regulator gene (Figure 1). Two contigs present in all ST131 encoded *araC* (a transcriptional regulator) and *sugE* (associated with resistance to quaternary ammonium salts). 39 AMR contigs in all bar 83972 included 28 copies of *mdfA*, encoding a multi-drug efflux pump linked to broad-spectrum AMR [82]. 3_2_53FAA and 83972 shared *ftsI* encoding a transpeptidase catalysing peptidoglycan crosslinking [83] with the synonym penicillin-binding protein 3 (PBP3) [84-85], and is sensitive to ESBLs [86].

For a wider context on ST131 dynamics, comparing this preterm infant resistome with *Klebsiella pneumoniae* showed that an interspecies shared resistome had 58 genes and that the larger *K. pneumoniae* core resistome of 308 genes included *bla*_SHV_ (Figure S1, Text S2). Nine of ten genes in pathogenic *K. pneumoniae* isolate PMK1 that were absent from a urinary tract microbiome assembly (WGLW1) were *bla*_OXY_, NCTC13441. Repeating this with the more divergent *Staphylococcus aureus* and *lugdunensis* found 14 shared AMR genes on *S. aureus* plasmids only (Figure S2, Text S3).

### Extensive plasmid rearrangements in closely related ST131 Clade C genomes

To evaluate the origin and relationships of the AMR genes in these seven Clade C (Table 1), we initially found with PlasmidFinder [94] that all had IncF1A and some C2 had lost IncF1B, suggesting potential plasmid changes (Table S3). We aligned these seven assemblies to plasmids common in ST131 (pEK499, pEK516, pEK204, pCA14, pV130) and repeated this via read mapping. Plasmids pEK499 (from England), pEK516 (England) and pEK204 (Belfast) were geographically near these ST131 (all from Ireland). SE15’s pECSF1 122.3 Kb IncF2A/F1B plasmid was used as a control because it had no known AMR genes [87]. 8289_1#35 and 8289_1#91 (both from C1) had identical results, and *vice versa* for 8289_1#60 and 8289_1#27 (both from C2) and so are not discussed below.

Clade C ST131 had more similarity to IncF2/F1A *bla*_CTX-M-15_-positive plasmid pEK499 than SE15 and the two HMP samples, but to varying degrees (Figure 2). This plasmid has 185 genes, lacks *traX* for conjugation, is stably inherited (Table S4), and may have been gained early in the origins of Clade C as a fusion of type F2 and F1A replicons [88]. Nine of pEK499’s ten key AMR genes are in a 25 Kb segment spanning *bla*_TEM_, *bla*_OXA-1_ and *bla*_CTX-M-15_ at 40, 58 and 63 Kb (respectively). 8289_1#3 and 8289_1#24 (both C2) had all three, 8289_1#27, 8289_1#60 and 8289_1#34 (all C2) had *bla*_OXA-1_ and *bla*_CTX-M-15_ but not *bla*_TEM_, whereas C1 (8289_1#35 and 8289_1#91) were *bla*_TEM_- and *bla*_CTX-M-14_-positive. Conjugation (*tra*) genes were in all ST131 except 8289_1#34. IncF2A plasmid pEK516 has ∼75% similarity to pEK499 but is shorter (64.6 Kb) [59] and is usually non-conjugative [89]. This plasmid has 103 genes including *bla*_TEM_, *bla*_OXA-1_ and *bla*_CTX-M-15_, and a type I partitioning locus for stable inheritance absent in pEK499 (Table S4). Read mapping to pEK516 showed that Clade C had high similarity to it, unlike SE15 and the HMP samples (Figure S3).

**Figure 2.**
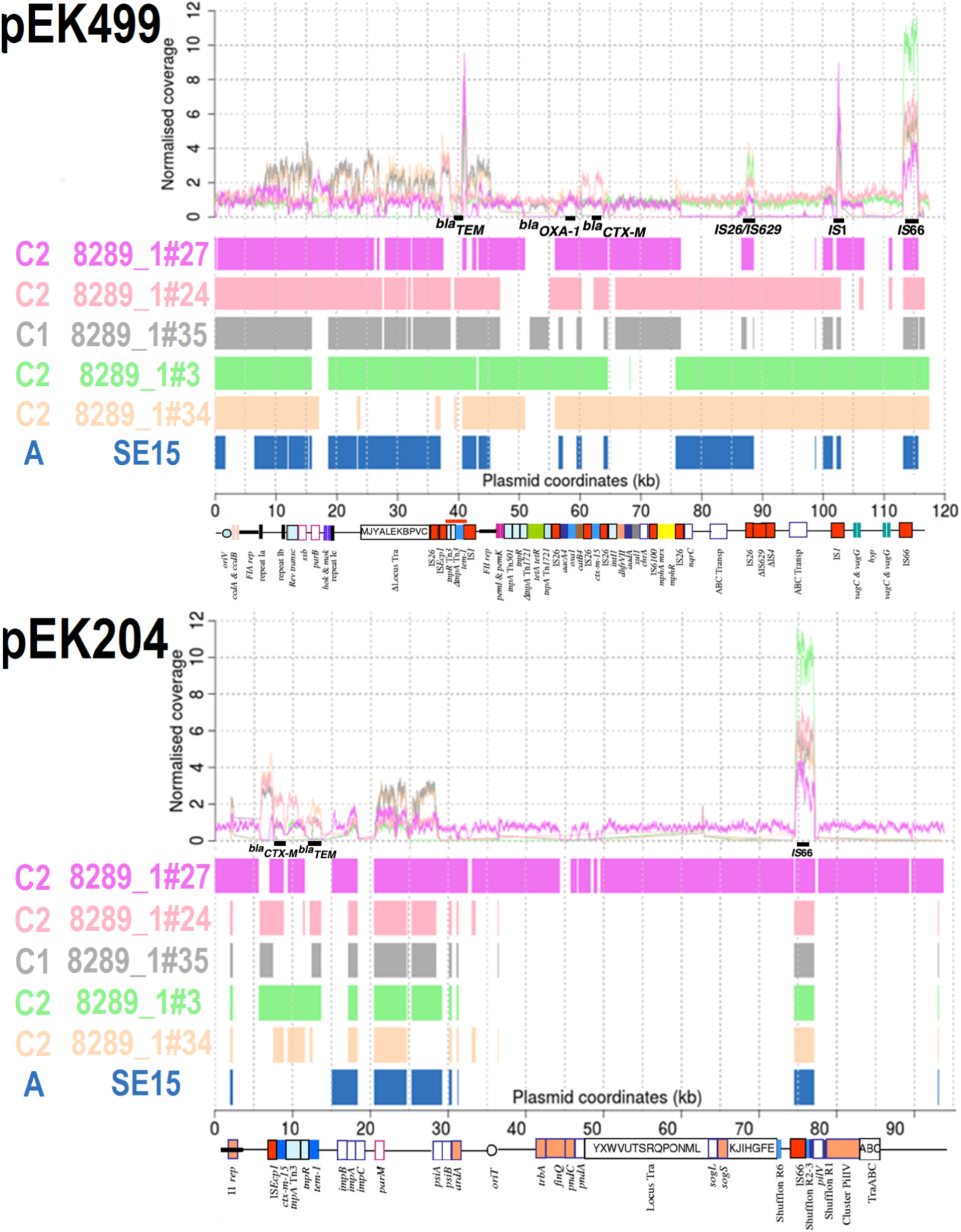
Comparison of pEK499 (top, 117,536 bp) and pEK204 (bottom, 93,732 bp) with five ST131 C1 and C2 isolates, ST131 Clade A reference SE15 (navy) and two HMP assemblies (3_2_53FAA in dark green and 83972 in cyan). Top: Normalised read coverage showed high copy numbers of IS*1* (at 41 and 103 Kb of pEK499) and IS*66* (at 75-77 Kb of pEK204, and at 113 Kb of pEK499). Bottom: BLAST alignments showed limited similarity for the HMP assemblies and SE15 relative to higher levels for Clade C for pEK499: 8289_1#27 (C2, mauve), 8289_1#24 (C2, pink), 8289_1#35 (C1, grey), 8289_1#3 (C2, light green), 8289_1#34 (C2, beige). For pEK204, only 8289_1#27 had many regions of similarity. Genes encoding *bla*_TEM_, *bla*_OXA-1_ and *bla*_CTX-M-15_ are at 40, 58 and 63 Kb on pEK499 (respectively). The *bla*_CTX-M_ gene was at 8 Kb, *bla*_TEM_ was at 13 Kb, followed by mixed conjugation and segregation genes at 36-70 Kb on pEK204. For pEK204, the *bla*_CTX-M-3_ gene differs from *bla*_CTX-M-15_ by a single R240G substitution, and so *bla*_CTX-M_ genes detected here were *bla*_CTX-M-15_. The annotation was modified from [59]. Matches spanning >300 bp are shown.

Contrasting sharply with pEK499 and pEK516, 8289_1#27 (C2) reads mapped to conjugative IncI1 plasmid pEK204 found 72.8 Kb regions of similarity, unlike the others that had <10 Kb (Figure 2). PEK204 is similar to IncI1 plasmid R64, has 112 genes, a complete *tra* region (Table S4), and a type I partitioning locus [59]. 8289_1#27 had a complete I1 replicon, *oriT, tra*, shufflons subject to site-specific recombinase activity, and a *pil* cluster encoding a type IV fimbriae associated with enhanced cell adherence and biofilm formation in entero-aggregative and Shiga toxin-producing *E. coli* [90]. A 9.3 Kb region on pEK204 contained *bla*_TEM-1b_, inactive transposase Tn*3-tnpA*, IS*Ecp1, bla*_CTX-M_ and a 5’ orf477-*tnpA-tnpR* region. The 14 bp IRL 5’ of IS*Ecp1* and IRR at the distal end of the inverted *orf477* element can mobilise *bla*_CTX-M_, and an additional IRR at *impB* 3’ of the *bla*_TEM-1b_ gene (7.4 Kb further away) can mobilise the 9.3 Kb unit [91], which has been found on IncFIA, IncFIA-FIB, IncN and IncY plasmids after originating on a pCOL1b-P9-like plasmid [59]. *ImpB* and *impA* encode an error-prone DNA repair protein (like UmuDC) [92].

Further investigation of the ST131 plasmid-matching regions showed variable plasmid similarity within closely related isolates. Read mapping to conjugative plasmid pCA14 showed that all Clade C bar 8289_1#3 had *Mrx* and *mph(A)* genes associated with macrolide resistance (Figure S4). This was also found by comparing with contigs from non-conjugative plasmids pV130a and pV130b from sewage treatment plant water in India [93], where 8289_1#27 and 8289_1#60 (both C2) had similarity spanning all the pV130a contigs (Figure S5).

### High rates of pEK499 and pEK204 protein-protein interactions with chromosomal *E. coli* proteins

The extensive diversity of Clade C AMR genes and plasmids raised the question of how the plasmids’ gene products interact with chromosomal ones. Protein-protein interaction (PPI) networks can examine plasmid-chromosome coevolution based on gene products’ topological proximity [95-96], which could be higher for plasmid and chromosomal proteins that have co-existed and so may interact more [97]. We used topological data analysis (TDA) to measure the number of non-trivial (indirect) loops where a loop is a chain of at least four PPIs ending at the same protein where it started. We focus on a type of loops called “non-trivial” (see Methods). The number of non-trivial loops was scaled by the number of PPIs per dataset (Table 2).

**Table 2.**
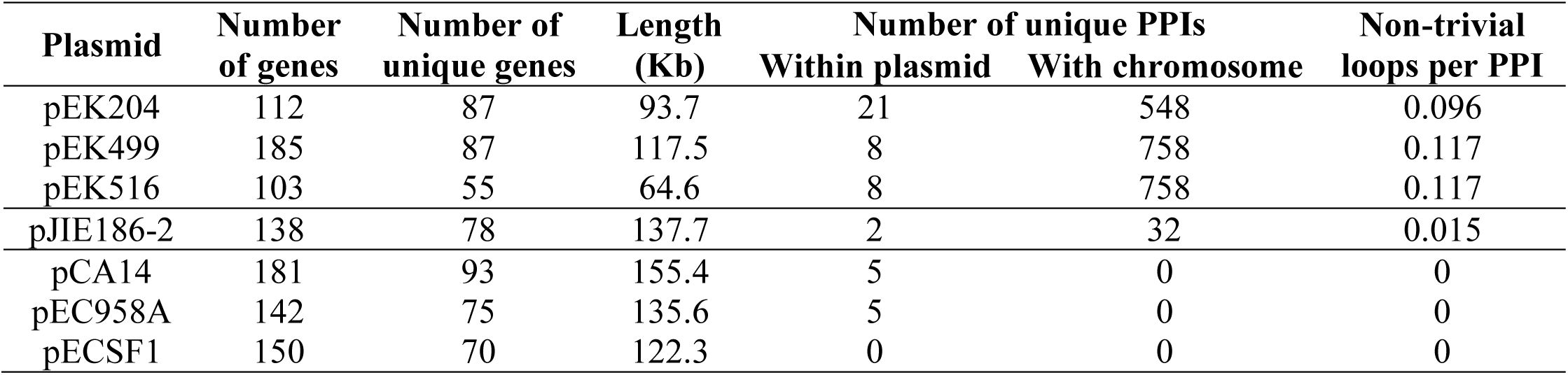
The numbers of protein-coding genes per plasmid, unique genes per plasmid, length, and numbers of non-redundant PPIs within the plasmid and with the chromosome, and the rate of indirect interactions (non-trivial loops) per PPI. Plasmid pEK516 had the same results as pEK499.

PPI data for 4,146 *E. coli* protein-coding genes with 105,457 PPIs (all combined scores >400) from the Search Tool for the Retrieval of Interacting Genes/Proteins (STRING) database v10 [98] was used to get the numbers of non-redundant PPIs and loops per PPI for a plasmid among its own proteins, and then between the plasmid’s proteins and the 4,146 chromosomal proteins. We assessed the numbers of PPIs from a combined score threshold of 400 to 900 with a step size of 25. We tested our approach using a published *E. coli* clustering of 60 genes [99] that had a rate of 0.244 non-trivial loops per PPI, which was lower than the rate of 0.518 obtained for the 4,146 chromosomal proteins (Table S5, Figure S6). There were lower rates of non-trivial loops per PPI for pEK499 (0.117) and pEK204 (0.096), but the plasmids’ rates were almost constant across the combined score thresholds (Figure S7). This held even though the number of PPIs per protein was negatively correlated with the combined score (Figure S8), suggesting that the rate of non-trivial loops per PPI was robust to changes in the combined score threshold. We found that pEK499 and pEK204 interacted with chromosomally encoded proteins, but pCA14, pEC958A and pECSF1 did not (Table S6), and pJIE186-2 had a small number of PPIs (Table 2) that may be VFs [100].

### ST131 genomes had an ancestral pEK499-like plasmid but some gained a pEK204-like one

The seven ST131 assemblies above were from a set of 4,071 ST131 from Clades A, B and C [61] that were aligned here to pEK499 and pEK204 with BLAST. 837 (20.6%) had >10 Kb of regions similar to pEK204 (Figure S9), whereas 3,108 (76.3%) had >10 Kb like pEK499 (Figure S10). All 193 assemblies with >40 Kb of pEK204-like segments were from Clade C: 17 from C0 (out of 52, 33%), 82 from C1 (out of 1,119, 7%) and 94 from C2 (out of 2,051, 5%) (Figure 3). All 17 from C0 had an I1 replicon, *bla*_CTX-M_ gene, *bla*_TEM_ gene (bar one isolate) and at least a partial *tra* region, and 13 had the *pil* operon (76%). The 82 from C1 and 94 from C2 had lower rates of I1 replicon presence (C1 77%, C2 81%), partial *tra* regions (C1 94%, C2 86%) and *pil* clusters (C1 n=60 or 73%, C2 n=71 or 76%) but differed in the rates of gene presence for *bla*_CTX-M_ (C1 28% vs C2 83%, odds ratio = 12.5, 95% CI 6.1-25.8, p=e-11) and *bla*_TEM_ (C1 59% vs C2 30%, odds ratio = 3.3, 95% CI 1.8-6.2, p=2.2e-5). This implied *bla*_CTX-M_ (via *orf477*) was common in C2 and *bla*_TEM_ (via *impB*) in C1 due to different ancestral transposition of the 9.3 Kb region.

**Figure 3.**
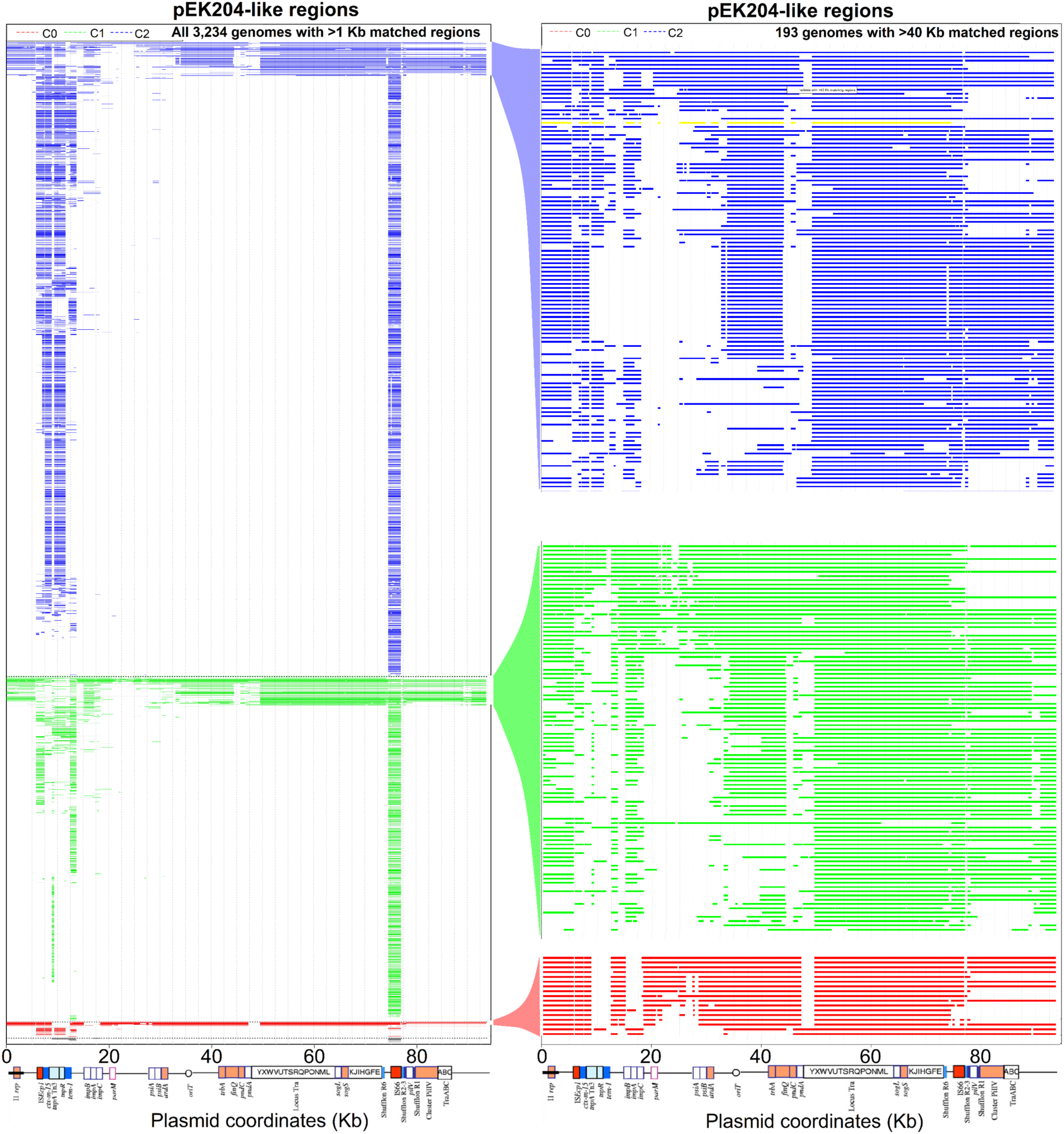
The distributions of pEK204-like regions (93,732 bp) in ST131 Clade C genome assemblies with >1 Kb (left, 3,234) and >40 Kb (right, 193) of matching segments. The regions of similarity were in (left) subclades B0 (14 in black), C0 (51 in red), C1 (1,119 in green), C2 (2,051 in blue); (right) C0 (17 in red), C1 (82 in green), C2 (94 in blue). 8289_1#27 (C2) is in yellow. Similarity across pEK204 was based on BLAST alignments. The *bla*_CTX-M_ (at 8 Kb) and *bla*_TEM_ (13 Kb) genes are followed by a mix of conjugation and segregation genes at 36-70 Kb. This suggested independent integrations and rearrangements of pEK204-like plasmids across subclades. The annotation was modified from [59].

To resolve the origin of pEK204-related *pil* HGT, we searched for the 14 *pil* operon genes (*pilIJKMNOPQRSTUVI)* individually in the 4,071 assemblies [61]. The entire *pil* operon was conserved in 376 (9%), including 61 from A (out of 414, 15%), 97 from B (out of 433, 22%), 17 from C0 (out of 52, 33%), 95 from C1 (out of 1,121, 8.5%) and 106 from C2 (out of 2,051, 5.2%) (Table 3) (see Data Access). Linked with the pEK204 matches, this indicated potential non-pEK204 *pil* ancestral acquisition in Clades A and B, and the putative Clade C ancestor may have had a pEK204-like plasmid with no *pil* that subsequently gained by recombination in some isolates.

**Table 3.** *Fim, pil, pap* and *ucl* operon genes detected in 4,071 ST131 genome assembles. The numbers and percentages of isolates per subclade with complete or absent (-) *pil* (*pilK*-*pilV*) or *pap* or *ucl* (*uclABCD*) operon genes where a subset also had three *pap* genes (*papIBA*). This focused on isolates allocated to the eleven main allelic options, isolates with some level of partial operon presence or absence were too sparse to be informative. *UclBCD* but not *uclA* was in an additional 2% (n=77), including 7 from A (2%), 28 from B (6%), none from C0, 24 from C1 (2%), and 18 from C2 (1%).

| Operon |  |  | Subclade |  |  |  |  |  |  |  |  |  | Totals |
| --- | --- | --- | --- | --- | --- | --- | --- | --- | --- | --- | --- | --- | --- |
| *pil* | *pap* | *ucl* | A | % | B | % | C0 | % | C1 | % | C2 | % |  |
| - | *papIBA* | - | 279 | 67% | 107 | 25% | 3 | 6% | 772 | 69% | 1,192 | 58% | 2,353 |
| - | all | all | 21 | 5% | 13 | 3% | 23 | 44% | 82 | 7% | 481 | 24% | 620 |
| all | *papIBA* | - | 57 | 14% | 2 | <1% |  |  | 75 | 7% | 66 | 3% | 200 |
| - | all | - | 1 | <1% | 94 | 22% |  |  | 8 | <1% | 14 | <1% | 117 |
| all | - | - | 3 | <1% | 71 | 16% |  |  | 5 | <1% | 3 | <1% | 82 |
| all | all | all |  |  | 16 | 4% | 1 | 2% | 10 | <1% | 33 | 2% | 60 |
| all | all | - | 1 | <1% | 6 | 1% | 16 | 31% |  |  | 4 | <1% | 27 |
| all | - | all |  |  |  |  |  |  | 1 | <1% |  |  | 1 |
| - | *papIBA* | all |  |  |  |  |  |  |  |  | 1 | <1% | 1 |
| all | *papIBA* | all |  |  |  |  |  |  |  |  |  |  | 0 |
| - | - | all |  |  |  |  |  |  |  |  |  |  | 0 |
| Partial matches |  |  | 52 | 13% | 124 | 29% | 9 | 17% | 168 | 15% | 257 | 13% | 610 |
| Totals |  |  | 414 |  | 433 |  | 52 |  | 1,121 |  | 2,051 |  | 4,071 |

We extended this to two other important fimbrial operons: *pap* (pyelonephritis-associated pili) and *ucl* (uroepithelial cell adhesin-like). All four *ucl* genes (*uclABCD*) were in 17.5% (712) of isolates, including 21 from A (5%), 30 from B (7%), 25 from C0 (48%), 94 from C1 (8%) and 542 from C2 (26%) (Table 3). All *ucl*-positive C2 genomes (bar one sample) had a complete *pap* operon. The *pap* operon was in a minority (26%, 1,049) of isolates, including 28 from A (7%), 163 from B (38%), 49 from C0 (94%), 127 from C1 (11%) and 682 from C2 (33%) (Table 3). Most isolates had *papIBA* but not the genes 3’ of them (*papHCDJKEF*) (65%, n=2,661): 359 from A (87%), 118 from B (27%), 3 from C0 (6%), 888 from C1 (79%) and 1,293 from C2 (63%). *PapI* (234 bp) is associated with higher P fimbriae expression in *E. coli* CFT073 [102] and here was amplified to two, three or four copies in 22% ST131: 26 in Clade A (6%), 51 in B (12%), 45 in C0 (87%), 119 in C1 (11%) and 658 in C2 (32%). Only 45 assemblies had an amplified *papB*, and 265 had an amplified *papA*. These two genes (*papBA*) share a promoter *pBA*, and *papI* has a separate one (*pI*).

Most isolates from A, C1 and C2 and a minority of B had no *pil* nor *ucl* operons and lost the *papH-F* segment (Table 3). Clade B differed from A because some from this diverse clade have either *pil* or *pap*, but not both. 83 ST131 with no detected *pap* genes all had a *pil* operon, and most (71 of 83) were from Clade B. Given the *fim* operon’s key role in ST131 evolution, we verified that *fim* was intact in >99% of the 4,071. As expected, the IS*Ec55* insertion at *fimB* [101] that delays *fim* expression in Clade C (like EC958 [20]) was in >99% of C1 (1,117, 99.6%) and C2 (2,041, 99.5%), but rarer in A (236, 57%), B (93, 21%) and C0 (18, 35%) (32 samples had additional *fimB* rearrangements). During the period prior to the divergence of C0 from C1 and C2 [50], the Clade C ancestor likely had a complete *pap* operon (but not *pil* nor *ucl*). The C1/C2 ancestor gained the *fimB* IS*Ec55* insertion and a minority of C1 gained either *pil* (80, 7%) or *ucl* (82, 7%) or both (11, 1%), whereas 24% of C2 gained *ucl* (482), 4% *pil* alone (73) and 2% both *ucl* and *pil* (33).

Unlike *ucl* gene products that are functionally independent [103], regulatory protein PapB reduces *fim* operon expression by inhibiting FimB and activating FimE, both tyrosine site-specific recombinases that invert *fimS*, including in *E. coli* CFT073 and 536 [104-106]. PapI and PapB regulate *pap* expression [105-107] depending on their concentrations and protein binding at the 416 bp regulatory region between *papI* and *papB*. Thus, retention of *papIBA* in >65% of ST131 could be linked to *fim* transcriptional regulation by PapB. Using the STRING PPI network data above for CFT073 and 536 that have *fim, pap* and *ucl* [108-109] (no *pil* gene products were found), there were ten *fim*-*pap* inter-operon PPIs in both 536 and CFT073, including pilus rod subunit PapA with FimD and FimF (as well as FimC), potentially matching the PPIs of FimA [110]. If only *papIBA* was present, the functional effects (if any) of *papI* and *papA* remain uncertain because it is unclear if FimC (replacing PapD at the inner cell membrane) could mediate periplasmic transport of PapA (like FimA) subunits via the chaperone FimI (instead of PapH) to usher FimD (rather than PapC) for pilus rod assembly at the outer cell membrane, extended later by FimF (in lieu of PapK) for the base of the tip. Nonetheless, presence of these genes can inform pilicide and coilicide design, such as antibodies targeting VF PapA [110].

## Discussion

AMR genes in commensal or environmental bacteria are a major reservoir for pathogens [111] driven by HGT and recombination in bacteria [112], including in *E. coli* [113-116], and microbiome species [117-118]. Previous work screened contigs from preterm infants for resistance to 16 antibiotics [74]. Here, we showed that this resistome was shared extensively between ExPEC ST131, commensal ST131 and microbiome *E. coli*, indicating likely transfer of these genes across commensal and pathogenic bacteria inhabiting the human gut and urinary tract, as expected [119].

*E. coli* 83972 does not express functional fimbrial adhesins and is used for therapeutic urinary bladder colonisation in patients in which it protects against super-infection [120-122]. Although 83972 lost virulence during its adaptation to commensalism [123], here it had AMR genes like PBP3 that it shared with an *E. coli* gut microbiome (3_2_53FAA). This retention of certain AMR genes in asymptomatic specimens is important when assessing the AMR gene evolution in ST131 [124-125].

Within ST131, Clade C core genomes are highly conserved but accessory genomes have extensive differences in AMR gene content [50,52,61]. Here, this was supported by NCTC13441’s repertoire of *bla*_TEM_ and *bla*_OXY_ genes, contrasting with EC958’s variety of *bla*_CMY_ ones. This reinforced the view that ST131’s accessory resistome is shaped by the environment mediated by plasmids, rather than population structure or geographic factors [126], so perhaps tracking plasmids and MGEs in addition to AMR genes [42,127] could assist treatment diagnostics [128].

By combining seven Clade C genome assemblies, then these with 4,064 Clade C assemblies, we found that most (78%) ST131 had >10 Kb of regions similar to plasmid pEK499 and some (9%) had regions with high similarity to IncI1 plasmid pEK204, suggesting either discrete gains of pEK204-like plasmids in each C subclade or its presence in the Clade C ancestral lineage. IncI1 plasmids are common in ExPEC and are associated with different ESBL genes [67,129-131]. Given that backbone plasmid genes may determine fitness effects more than ESBL genes [132] and long-term IncF plasmid persistence in ST131 [62], plasmids pEK204, (pEK516,) and pEK499 proteins’ higher interaction rates with chromosomally-encoded proteins relative to other relevant plasmids could be due to co-evolution. Measuring the indirect connectivity as PPI network loops per interaction showed evidence of this long-term retention, and may help identify plasmids compatible with *E. coli* chromosomes.

ST131’s fitness advantage is tightly correlated with type 1 fimbriae variant *fimH30* and delayed *fim* operon expression due to an insertion at *fimB* [16,133-134]. Here, a minority of ST131 had type IV pilus biosynthesis (*pil*) genes and most Clade C with *pil* had >40 Kb of regions similar to pEK204, whereas Clade A and B isolates with *pil* did not have pEK204-like regions. *Pil* may allow different epithelial cell adhesion from *fim* via a thinner pilus and biofilm formation [90]. Additionally, F17-like *ucl* operon was more common in C2 (24%): UclD (like FimH) is a two-domain tip adhesin that binds intestinal epithelial cells via O-glycans (FimH uses N-glycans) [103]. The *ucl* operon is associated with uroepithelial cell adhesion, biofilm formation and could be mobilised by its flanking MGEs [135].

The *pap* operon encodes a P fimbriae with high specificity for kidney epithelial cell and erythrocyte receptor glycolipids (α-D-galactopyranosol(1-4)-α-D-galactopyranoside) [136]. Delayed *fim* expression in Clade C [20,101] may be reduced further by PapB: *papIBA* alone was retained in >65% of Clade C here. Delayed *fim* expression [137] and having multiple P fimbrial gene clusters [107] associates more with pyleonephritis than cystitis. Given that *E. coli* express single fimbriae at the cell surface [104,138] and isogenic cell populations can express distinct fimbrial types [139], type 1 fimbriae may allow bladder colonisation followed by reduced expression in favour of P fimbriae when at the kidney [138]. Our results on plasmid-linked changes across the ST131 resistome and diversity of fimbrial gene composition could inform on potential infection [140] and biofilm formation [20].

## Methods

### *E. coli* genome isolate collection

Of the 12 *E. coli* assemblies examined (Table 1), three were ST131 references: SE15, NCTC13441 and EC958. Seven were ST131 from Ireland in 2005-2010, of which two from C1 were *bla*_CTX-M-14_-positive, as was 8289_1#24 from C2, and all five from C2 were *bla*_CTX-M-15_-positive. All seven were from urine except 8289_1#34, which was a rectal swab. The two HMP samples were: 3_2_53FAA (aka EC3_2_53FAA) from a 52-year-old male Canadian with Crohn’s disease’s colon biopsy [141]. 83972 (aka EC83972) was from the urine of a Swedish girl with a three-year history of asymptomatic bacteriuria and stable renal function [142]. 83972’s has common ancestry with virulent pyelonephritis-causing ExPEC CFT073 [143]. ST131 C2 reference genomes NCTC13441 and EC958 were from UTIs in the UK, and had *bla*_CTX-M-15_-positive plasmids [9,59,133]. NCTC13441 has 4,983 predicted protein-coding genes and EC958 has 4,982. EC958 has numerous virulence-associated genes that encode adhesins, autotransporter proteins and siderophore receptors, and can cause impairment of uterine contractility in mice [144]. Plasmid pEC958A has 85% similarity with pEK499 but lacks the latter’s second *tra* region due to an IS26-mediated *bla*_TEM-1_ insertion [9,60,133]. ST131 Clade A reference SE15 was examined as a genetic outgroup and a commensal control because it lacks many virulence-associated genes. It has a 4,717,338 bp chromosome with 4,338 protein-coding genes and a 122 Kb plasmid pSE15 with 150 protein-coding genes [87].

### Illumina library quality control and read mapping

The paired-end Illumina HiSeq libraries for each sample were screened for low quality (Phred score <30) and short (<50 bp) reads using Trimmomatic v0.36 [145] and corrected using BayesHammer from SPAdes v3.9 [145] (Table S7). These corrected read libraries were mapped to references with SMALT v7.6 (www.sanger.ac.uk/resources/software/smalt/), and the SAM files were converted to BAM format, sorted and PCR duplicates removed using SAMtools v1.19 [146].

### Homology-based resistome screening and comparison

The read mapping to the reference resistome (Table S1, see Methods of [74]) used GROOT [78] where the reads were indexed using the median read length (Table S7). Contig and protein domain annotation was derived from the Pfam v27.0 and ProSite databases using InterProScan v5.22-61 [147]. The protein homolog dataset in the CARD (2,239 genes) [77] was aligned with the genomes to annotate the resistomes with BLAST v2.2.31, where matches with a bit score >500 and >99% homology were considered valid. Alignments were visualised using Artemis and the Artemis Comparison Tool (ACT) [148], and also R packages VennDiagram v1.6.1, Seqinr v3.4-5, UpSetR v1.4.0 and WriteXLS v5.0.0.

### Assessment of plasmids prevalent in ST131

Sequence and annotation files for pEK499 (NC_013122.1, EU935739), pEK516 (NC_013121.1, EU935738), pEK204 (NC_013120.1, EU935740) and all pV130 contigs (LC056314.1 to LC056328.1) [93] were aligned with Clustal Omega v1.2.1 [149]. Replicon typing used PlasmidFinder [56] and each plasmid was compared to the CARD. Each read library was mapped to each plasmid to verify local genetic features and quantify copy number levels, visualised with Artemis and R v3.4.2’s Reshape2 v1.4.3, Ggridges v0.5.1, Ggplot2 v2.3.2.1, Readr v1.3.1, Dplyr v0.8.3 and Ape v5.3 packages. Sequence similarity was calculated using the Sequence Identity and Similarity (SIAS) tool (http://imed.med.ucm.es/Tools/sias.html). Homology searches against pEK204 and pEK499 in the 4,071 genomes [60] examined matches >300 bp length with >98% sequence identity.

### Protein-protein interaction network construction and topological data analysis

TDA has been used to investigate complex and high-dimensional datasets, like breast cancer genomes [150]. We applied it to PPI networks with a focus on direct and indirect connectivity across different network topologies by quantifying rates of absent PPIs (non-trivial loops). The numbers of non-trivial loops per PPI was used as a metric for indirect connectivity because it was consistent across parameters and was independent of network size. We assessed pCA14, pEK204, pEK499, pEK516 and pEC958A, along with two plasmids with no known AMR genes as negative controls: pECSF1 and pJIE186-2 [100]. We examined unique genes with PPI network information only. Results for pEK516 were identical to those for pEK499.

We extracted *E. coli* K12 MG1655 PPI data from the STRING database v10 [98] with R v3.5.2 packages BiocManager v1.30.4, Dplyr v0.8.0.1, Genbankr v1.10.0, Rentrez v1.2.1, STRINGdb v1.22.0, tidyverse v1.2.1 and VennDiagram v1.6.20. STRING’s combined scores were used because they integrate multiple types of evidence while controlling for random PPIs [98]. Using a score threshold of 400, K12 had 4,146 protein-coding genes, of which 4,121 had interactions, resulting in 105,457 PPIs – 14,555 PPIs were present for a score threshold of 900. For each plasmid or set of genes, we obtained the unique genes and numbers of pairwise PPIs within that set and with the chromosome. We tested our TDA-based approach using previous work [99]. For *E. coli* 536, FooB was used as an equivalent for PapB, and likewise for F7-2 that matched VF PapA in CFT073, and FooG for PapG.

We constructed a Vietoris-Rips complex [151] where the proteins were the vertices with PPIs as edges, so that proteins A and B with a PPI would be joined by a single edge (two proteins in one dimension, 1-D). Proteins A, B and C joined by three PPIs would have a filled 2-D triangle. For four proteins A, B, C and D joined by all six possible pairwise PPIs, their four filled 2-D triangles A/B/C, B/C/D, A/C/D and A/B/D constitute a tetrahedral surface filled with a 3-D tetrahedron. This can be extended to *m+1* pairwise connected proteins, which get a *m-D* polytope filled into their skeleton of edges, triangles, tetrahedra, and so on. For *m≥4*, a loop connects *m* proteins by *m* PPIs, such that each protein is involved in precisely two of these PPIs. For instance, four proteins A, B, C, D can be joined to a loop by four PPIs A-B, B-C, C-D, D-A, where A-B is a PPI between the proteins A and B (etc). Any loop that can be filled with triangles, along existing PPIs, is called trivial. So if there is an interaction between A and C, the loop is filled by the two triangles AB-BC-CA and CD-DA-CA and hence trivial. If there are no interactions between A and C, and none between B and D, then the loop is non-trivial.

Specialised software [152] was used to optimise the Betti number computations instead of using general purpose implementations of the Vietoris-Rips complex (such as SAGE) due to the large K12 dataset size (4,146 proteins with 105,457 interactions). This software used the sparsity of the boundary matrices to process the rank computations efficiently in LinBox [153], with a hard-coded dimensional truncation of the Vietoris-Rips complex to avoid the large number of high-dimension simplices that would be obtained for the full dataset. For each analysis across STRING combined scores 400 to 900 with a step of 25, we computed the first Betti number of the Vietoris-Rips complex that counted the numbers of non-trivial loops (with missing PPIs inside), which was adjusted for the numbers of PPIs above the score threshold (i.e., loops per PPI). The number of PPIs for the complete K12 chromosomal dataset was negatively correlated with the combined score threshold (r^2^=0.964), as was the loops per PPI (r^2^=0.905), so the chromosomal loops per PPI was used as a baseline for different score thresholds.

### Operon gene homology search approach

Homology searches for each *fim* operon gene were implemented using the NCTC13441 annotation coordinates to extract the corresponding sequence with SAMtools v1.9 and align each gene to the 4,071 ST131 assemblies [60] with BLAST v2.2.31, processed with R packages Tidyverse v1.2.1 and Dplyr v0.8.3. For all operons, minor individual gene partial matches or losses were not examined due to the large number of samples and lack of consistent patterns for rare combinations. Results for amplified genes were restricted to those with prevalence >1%. The CDS of *fimB* is 600 bp, and with the IS*Ec55* insertion the region typically spanned 2,493 bp. NCTC13441 has only three *pap* genes (*papIBA*) and lacks any *ucl* or *pil* genes, so the *pil* genes were from pEK204. *PilI* and *pilJ* encoding IncI1 conjugal transfer proteins were mainly absent with no clear association with the other *pil* genes, and so were not examined here. *PilV* encoding a pilus tip adhesin had >1 copy in 292 assemblies, as expected for a locus rearranged to change pilus binding specificity [154]. The sequences for the four genes in the 5 Kb *ucl* operon were determined from *E. coli* 83972 (CP001671): major subunit (*uclA*), chaperone (*uclB*), usher (*uclC*) and adhesin (*uclD*). *E. coli* UTI89’s chromosome (CP000243) and plasmid (CP000244) [112] were used to get the *pap* genes.

## Supplementary Data

Table S1 – The 16 antibiotics used to identify 794 AMR genes on contigs originally from preterm infants.

Table S2 – The five AMR genes unique to all ST131 Clade C genomes except 8289_1#27 encoded *bla*_TEM_ genes (an Ambler class A β-lactamase).

Text S1 – The preterm infant resistome in ST131 genomes including NCTC13441.

Text S2 – A small shared infant resistome between ST131 and *Klebsiella pneumoniae*

Figure S1 – Chromosomal AMR genes in *K. pneumoniae* reference isolate PMK1 versus a *K. pneumoniae* microbiome (WGLW1), relative to *E. coli* ST131 EC958 and commensal *E. coli* SE15.

Text S3 – No shared infant resistome between ST131 and *Staphylococcus lugdunensis*

Figure S2 – Out of 794 preterm infant AMR genes, there were no overlapping chromosomal genes in *S. lugdenensis* reference HKU versus *S. lugdenensis* microbiome (M23590) relative to ST131.

Table S3 – PlasmidFinder Inc group alignments against pV130 and seven ST131 isolates.

Table S4 – Key known AMR, plasmid persistence and conjugation genes for pEK204, pEK499, pEK516, pCA14, pV130a and pV130b.

Figure S3 – Comparison of pEK516 (64,471 bp) with ST131 and two HMP assemblies.

Figure S4 – Read mapping distributions for seven ST131 to pV130 contigs.

Figure S5 – Comparison of plasmid pCA14 (155 Kb) with seven ST131 C1/C2 isolates.

Table S5 – The PPIN characteristics of the protein-coding genes from Miryala et al (2018).

Table S6 – The seven plasmids’ accessions, numbers of genes with interactions given a combined score of 400+, Inc groups, conjugative ability and AMR genes.

Figure S6 – The ratio of non-trivial loops per PPI (y-axis) versus the STRING combined score for all 60 genes from Miryala et al (2018) Table 1 [99].

Figure S7 – The ratio of non-trivial loops per interaction versus the STRING combined score for all chromosomal genes, all genes on pEK499 and all those on pEK204.

Figure S8 – The ratio of interactions per protein for genes encoded on the chromosome or the pEK499 and pEK204 plasmids across the STRING combined score.

Figure S9 – The distributions of pEK204-like and pEK499-like regions in ST131 assemblies.

Figure S10 – The distributions of pEK204-like regions in ST131 Clade C genome assemblies.

Table S7 – Read library summary statistics for each ST131 sample and reference genome.

## Author Contributions

Conceptualisation, AD, AR, TD; Software, Validation, Investigation and Visualisation, AD, AR, HA, BA, LC, KE, LM, MN, NT, SP, GS, CS, ZV, CW; Methodology, AD, AD, NT, TD; Writing – Original Draft Preparation, AD, NT, AR, TD; Writing – Review & Editing, AD, AR, TD; Supervision and Project Administration, AD, AR, TD. Funding, AD, TD.

## References

1. Ben Zakour NL, et al. Sequential acquisition of virulence and fluoroquinolone resistance has shaped the evolution of *Escherichia coli* ST131. mBio 2016 7, e00347, https://doi.org/10.1128/mBio.00347-16.

2. Calhau V, Ribeiro G, Mendonça N, Da Silva GJ. Prevalent combination of virulence and plasmidic-encoded resistance in ST 131 *Escherichia coli* strains. Virulence 2013 4(8), 726–9, https://doi.org/10.4161/viru.26552.

3. Van der Bij AK, Peirano, G, Pitondo-Silva A, Pitout JD. The presence of genes encoding for different virulence factors in clonally related *Escherichia coli* that produce CTX-Ms. Diagn Microbiol Infect Dis 2012 72, 297–302, https://doi.org/10.1016/j.diagmicrobio.2011.12.011.

4. Goswami C, Fox S, Holden M, Connor M, Leanord A, Evans TJ. Genetic analysis of invasive *Escherichia coli* in Scotland reveals determinants of healthcare-associated versus community-acquired infections. Microb Genom. 2018 4(6), doi: 10.1099/mgen.0.000190

5. Poolman JT, Wacker, M. Extraintestinal Pathogenic *Escherichia coli*, a Common Human Pathogen: Challenges for Vaccine Development and Progress in the Field. J Infect Dis. 2016 213(1), 6–13, https://doi.org/10.1093/infdis/jiv429.

6. Livermore D. The 2018 Garrod Lecture: Preparing for the Black Swans of resistance. Journal of Antimicrobial Chemotherapy 2018 73, 2907–2915, https://doi.org/10.1093/jac/dky265

7. Banerjee R, Johnson JR. A new clone sweeps clean: the enigmatic emergence of *Escherichia coli* sequence type 131. Antimicrob Agents Chemother 2014 58, 4997–5004, https://doi.org/10.1128/AAC.02824-14.

8. Pitout JDD, DeVinney R. *Escherichia coli* ST131: a multidrug-resistant clone primed for global domination. F1000 Research 2017 6(F1000 Faculty Review), 195.

9. Forde BM, et al. The complete genome sequence of *Escherichia coli* EC958: A high quality reference sequence for the globally disseminated multidrug resistant *E. coli* O25b:H4-ST131 clone. Plos One 2014 9(8), pp.e104400.

10. McNally A, et al. Combined analysis of variation in core, accessory and regulatory genome regions provides a super-resolution view into the evolution of bacterial populations. PLoS Genet 2016 12, 1006280.

11. Kuehn MJ, Heuser J, Normark S, Hultgren SJ. P pili in uropathogenic *E. coli* are composite fibres with distinct fibrillar adhesive tips. Nature 1992 356(6366), 252–5

12. Connell I, Agace W, Klemm P, Schembri M, Marild S, Svanborg C. Type 1 fimbrial expression enhances *Escherichia coli* virulence for the urinary tract. Proc Natl Acad Sci U S A 1996 93(18), 9827–321996.

13. Martinez JJ, Mulvey MA, Schilling JD, Pinkner JS, Hultgren SJ. Type 1 pilus-mediated bacterial invasion of bladder epithelial cells. EMBO J. 2000 19(12), 2803–122000

14. Selvarangan R, Goluszko P, Singhal J, Carnoy C, Moseley S, Hudson B, Nowicki S, Nowicki B. Interaction of Dr adhesin with collagen type IV is a critical step in *Escherichia coli* renal persistence. Infect Immun. 2004 72(8), 4827–352004.

15. Price LB, et al. The epidemic of extended-spectrum-beta-lactamase-producing *Escherichia coli* ST131 is driven by a single highly pathogenic subclone, H30-Rx. mBio 2013 4, e00377–13, https://doi.org/10.1128/mBio.00377-13.

16. Petty NK, et al. Global dissemination of a multidrug resistant Escherichia coli clone. Proc Natl Acad Sci U S A. 2014 111(15), 5694–9. doi: 10.1073/pnas.1322678111

17. Bloch CA, Stocker BA, Orndorff PE. A key role for type 1 pili in enterobacterial communicability. Mol. Microbiol. 1992 6, 697–701.

18. Bloch CA, Orndorff PE. Impaired colonization by and full invasiveness of *Escherichia coli* K1 bearing a site-directed mutation in the type1 pilin gene. Infect. Immun. 1990 58, 275–278

19. Totsika M, et al. A FimH inhibitor prevents acute bladder infection and treats chronic cystitis caused by multidrug-resistant uropathogenic *Escherichia coli* ST131. J Infect Dis 2013 208, 921–8.

20. Sarkar S, Vagenas D, Schembri MA, Totsika M. Biofilm formation by multidrug resistant *Escherichia coli* ST131 is dependent on type 1 fimbriae and assay conditions. Pathog Dis 2016 74 doi: 10.1093/femspd/ftw013.

21. Bahrani-Mougeot FK, Buckles EL, Lockatell CV, Hebel JR, Johnson DE, Tang CM, Donnenberg MS. Type 1 fimbriae and extracellular polysaccharides are preeminent uropathogenic *Escherichia coli* virulence determinants in the murine urinary tract. Mol. Microbiol. 2002 45, 1079–1093.

22. Bäumler AJ, Sperandio V. Interactions between the microbiota and pathogenic bacteria in the gut. Nature 2016 535(7610), 85–93. doi: 10.1038/nature18849

23. Bittinger K, et al. Bacterial colonization reprograms the neonatal gut metabolome. Nat Microbiol 2020 doi: 10.1038/s41564-020-0694-0

24. Conway T, Cohen PS. Commensal and Pathogenic *Escherichia coli* Metabolism in the Gut. Microbiol Spectr. 2015 3(3) doi: 10.1128/microbiolspec.MBP-0006-2014

25. Hudault S, Guignot J, Servin AL. *Escherichia coli* strains colonising the gastrointestinal tract protect germfree mice against Salmonella typhimurium infection. Gut 2001 49(1), 47–55

26. Li G, et al. Towards understanding global patterns of antimicrobial use and resistance in neonatal sepsis: insights from the NeoAMR network. Arch Dis Child. 2020 105(1), 26–31. doi: 10.1136/archdischild-2019-316816

27. Groussin M, et al. Industrialization is associated with elevated rates of horizontal gene transfer in the human microbiome. BioRxiv 2020 doi:http://dx.doi.org/10.1101/2020.01.28.922104

28. Gurnee EA, et al. Gut Colonization of Healthy Children and Their Mothers With Pathogenic Ciprofloxacin-Resistant *Escherichia coli*. J Infect Dis. 2015 212(12), 1862–8. doi: 10.1093/infdis/jiv278

29. Yamamoto S, Tsukamoto T, Terai A, Kurazono H, Takeda Y, Yoshida O. Genetic evidence supporting the fecal-perineal-urethral hypothesis in cystitis caused by *Escherichia coli*. J Urol 1997 157, 1127–9.

30. Rodríguez-Revuelta MJ, López-Cerero L, Serrano L, Luna-Lagares S, Pascual A, Rodríguez-Baño J. Duration of Colonization by Extended-Spectrum β-Lactamase-Producing Enterobacteriaceae in Healthy Newborns and Associated Risk Factors: A Prospective Cohort Study. Open Forum Infect Dis. 2018 5(12), ofy312 doi: 10.1093/ofid/ofy312

31. Roemhild R, Linkevicius M, Andersson DI. Molecular mechanisms of collateral sensitivity to the antibiotic nitrofurantoin. PLoS Biol. 2020 18(1), e3000612. doi: 10.1371/journal.pbio.3000612

32. Russell JT, et al. Antibiotics may influence gut microbiome signaling to the brain in preterm neonates. BioRxiv 2020 doi:https://doi.org/10.1101/2020.04.20.052142

33. ECDC, European Centre for Disease Prevention and Control. European Centre for Disease Prevention and Control. Antimicrobial resistance surveillance in Europe 2015. Annual Report of the European Antimicrobial Resistance Surveillance Network (EARS-Net). 2017 Stockholm: ECDC.

34. Findlay J, Gould VC, North P, Bowker KE, Williams MO, MacGowan AP, Avison MB. Characterization of cefotaxime-resistant urinary *Escherichia coli* from primary care in South-West England 2017-18. J Antimicrob Chemother. 2020 75(1), 65–71. doi: 10.1093/jac/dkz397

35. Mathers AJ, Peirano G, Pitout JD. The role of epidemic resistance plasmids and international high-risk clones in the spread of multidrug-resistant *Enterobacteriaceae*. Clin Microbiol Rev. 2015 28(3), 565–91, https://doi.org/10.1128/CMR.00116-14

36. Pitout JD, Laupland KB. Extended-spectrum beta-lactamase-producing Enterobacteriaceae: an emerging public-health concern. Lancet Infect Dis. 2008 8(3), 159–66. doi: 10.1016/S1473-3099(08)70041-0

37. Stokes HW, Gillings MR. Gene flow, mobile genetic elements and the recruitment of antibiotic resistance genes into Gram-negative pathogens. FEMS Microbiol Rev 2011 35, 790–819, doi: 10.1111/j.1574-6976.2011.00273.x

38. Von Wintersdorff CJ, et al. Dissemination of Antimicrobial Resistance in Microbial Ecosystems through Horizontal Gene Transfer. Front Microbiol 2016 7, 173, doi: 10.3389/fmicb.2016.00173.

39. Goldstone RJ, Smith DGE. A population genomics approach to exploiting the accessory ‘resistome’ of *Escherichia coli*. Microb Genom. 2017 3(4), e000108. doi: 10.1099/mgen.0.000108

40. Baraniak A, Fiett J, Hryniewicz W, Nordmann, Gniadkowski M. Ceftazidime-hydrolysing CTX-M-15 extended-spectrum beta-lactamase (ESBL) in Poland. J Antimicrob Chemother. 2002 50(3), 393–6

41. Pehrsson EC, Tsukayama P, Patel S, Mejía-Bautista M, Sosa-Soto G, Navarrete KM, Calderon M, Cabrera L, Hoyos-Arango W, Bertoli MT, Berg DE, Gilman RH, Dantas G. Interconnected microbiomes and resistomes in low-income human habitats. Nature. 2016 533(7602), 212–6. doi: 10.1038/nature17672

42. Durrant MG, Li MM, Siranosian BA, Montgomery SB, Bhatt AS. A Bioinformatic Analysis of Integrative Mobile Genetic Elements Highlights Their Role in Bacterial Adaptation. Cell Host Microbe. 2020 27(1), 140–153.e9. doi: 10.1016/j.chom.2019.10.022

43. Cerqueira GC, et al. Multi-institute analysis of carbapenem resistance reveals remarkable diversity, unexplained mechanisms, and limited clonal outbreaks. Proc Natl Acad Sci U S A. 2017 114(5), 1135–1140. doi: 10.1073/pnas.1616248114

44. Hazen TH, et al. Diversity among bla(KPC)-containing plasmids in *Escherichia coli* and other bacterial species isolated from the same patients. Sci Rep. 2018 8(1), 10291. doi: 10.1038/s41598-018-28085-7

45. Kwong JC, et al. Translating genomics into practice for real-time surveillance and response to carbapenemase-producing *Enterobacteriaceae*: evidence from a complex multi-institutional KPC outbreak. PeerJ 2018 6, e4210. doi: 10.7717/peerj.4210

46. Bosch T, et al. Outbreak of NDM-1-Producing *Klebsiella pneumoniae* in a Dutch Hospital, with Interspecies Transfer of the Resistance Plasmid and Unexpected Occurrence in Unrelated Health Care Centers. J Clin Microbiol. 2017 55(8), 2380–2390. doi: 10.1128/JCM.00535-17

47. Jamrozy D, et al. Evolution of mobile genetic element composition in an epidemic methicillin-resistant *Staphylococcus aureus*: temporal changes correlated with frequent loss and gain events. BMC Genomics 2017 18(1), 684. doi: 10.1186/s12864-017-4065-z

48. Martin J, et al. Covert dissemination of carbapenemase-producing *Klebsiella pneumoniae* (KPC) in a successfully controlled outbreak: long- and short-read whole-genome sequencing demonstrate multiple genetic modes of transmission. J Antimicrob Chemother. 2017 72(11), 3025–3034. doi: 10.1093/jac/dkx264

49. Smet A, Rasschaert G, Martel A, Persoons D, Dewulf J, Butaye P, Catry B, Haesebrouck F, Herman L, Heyndrickx M. *In situ* ESBL conjugation from avian to human *Escherichia coli* during cefotaxime administration. J Appl Microbiol. 2011 110(2), 541–9. doi: 10.1111/j.1365-2672.2010.04907.x

50. Ludden C, Decano AG, Jamrozy D, Pickard D, Morris D, Parkhill J, Peacock SJ, Cormican M, Downing T. Genomic surveillance of *Escherichia coli* ST131 identifies local expansion and serial replacement of subclones. Microb Genom. 2020 doi: 10.1099/mgen.0.000352

51. McNally A, et al. Diversification of Colonization Factors in a Multidrug-Resistant *Escherichia coli* Lineage Evolving under Negative Frequency-Dependent Selection. MBio. 2019 10(2), e00644–19, https://doi.org/10.1128/mBio.00644-19.

52. Decano AG, Ludden C, Feltwell T, Judge K, Parkhill J, Downing T. Complete assembly of *Escherichia coli* ST131 genomes using long reads demonstrates antibiotic resistance gene variation within diverse plasmid and chromosomal contexts. mSphere 2019 4(3), e00130–19, https://doi.org/10.1128/mSphere.00130-19.

53. Ny S, Sandegren L, Salemi M, Giske CG. Genome and plasmid diversity of Extended-Spectrum beta-Lactamase-producing *Escherichia coli* ST131 – tracking phylogenetic trajectories with Bayesian inference. Scientific Reports 2019 9, 10291, https://doi.org/10.1038/s41598-019-46580-3.

54. Kallonen T, et al. Systematic longitudinal survey of invasive *Escherichia coli* in England demonstrates a stable population structure only transiently disturbed by the emergence of ST131. Genome Res. 2017 27, 1437–1449, https://doi.org/10.1101/gr.216606.116.

55. Johnson JR, et al. Household Clustering of *Escherichia coli* Sequence Type 131 Clinical and Fecal Isolates According to Whole Genome Sequence Analysis. Open Forum Infect Dis. 2016 3(3), ofw129.

56. Carattoli A. Resistance plasmid families in *Enterobacteriaceae*. Antimicrob Agents Chemother. 2009 53(6), 2227–38. doi: 10.1128/AAC.01707-08

57. Nicolas-Chanoine MH, et al. Intercontinental emergence of *Escherichia coli* clone O25:H4-ST131 producing CTX-M-15. J Antimicrob Chemother. 2008 61(2), 273–81.

58. Nicolas-Chanoine MH, Bertrand X, Madec JY. *Escherichia coli* ST131, an intriguing clonal group. Clin Microbiol Rev. 2014 27(3), 543–74. doi: 10.1128/CMR.00125-13

59. Woodford N, Carattoli A, Karisik E, Underwood A, Ellington MJ, Livermore DM. Complete nucleotide sequences of plasmids pEK204, pEK499, and pEK516, encoding CTX-M enzymes in three major *Escherichia coli* lineages from the United Kingdom, all belonging to the international O25:H4-ST131 clone. Antimicrob. Agents Chemother. 2009 53(10), 4472–4482.

60. Phan MD, et al. Molecular characterization of a multidrug resistance IncF plasmid from the globally disseminated *Escherichia coli* ST131 clone. PLoS One. 2015 10(4), e0122369. doi: 10.1371/journal.pone.0122369

61. Decano AG, Downing T. An *Escherichia coli* ST131 pangenome atlas reveals population structure and evolution across 4,071 isolates. Sci Rep. 2019 9(1), 17394. doi: 10.1038/s41598-019-54004-5

62. Goswami C, Fox S, Holden MTG, Connor M, Leanord A, Evans TJ. Origin, maintenance and spread of antibiotic resistance genes within plasmids and chromosomes of bloodstream isolates of *Escherichia coli*. Microb Genom. 2020 doi: 10.1099/mgen.0.000353

63. Jørgensen SB, Søraas A, Sundsfjord A, Liestøl K, Leegaard TM, Jenum PA. Fecal carriage of extended spectrum β-lactamase producing *Escherichia coli* and *Klebsiella pneumoniae* after urinary tract infection – A three year prospective cohort study. PLoS One. 2017 12(3):e0173510. doi: 10.1371/journal.pone.0173510

64. Coyne MJ, Zitomersky NL, McGuire AM, Earl AM, Comstock LE. Evidence of extensive DNA transfer between bacteroidales species within the human gut. MBio 2014 5, e01305–14.

65. Zhao S, et al. Adaptive evolution within the gut microbiome of individual people. BioRxiv 2017 doi: 10.1101/208009

66. Bishara A, et al. Strain-resolved microbiome sequencing reveals mobile elements that drive bacterial competition on a clinical timescale. BioRxiv 2017 doi: 10.1101/125211

67. Knudsen PK, et al. Transfer of a *bla*CTX-M-1-carrying plasmid between different *Escherichia coli* strains within the human gut explored by whole genome sequencing analyses. Sci Rep. 2018 8(1), 280. doi: 10.1038/s41598-017-18659-2

68. Göttig S, Gruber TM, Stecher B, Wichelhaus TA, Kempf VA. In vivo horizontal gene transfer of the carbapenemase OXA-48 during a nosocomial outbreak. Clin Infect Dis. 2015 60(12), 1808–15. doi: 10.1093/cid/civ191

69. Willemsen I, van Esser J, Kluytmans-van den Bergh M, Zhou K, Rossen JW, Verhulst C, Verduin K, Kluytmans J. Retrospective identification of a previously undetected clinical case of OXA-48-producing *K. pneumoniae* and *E. coli*: the importance of adequate detection guidelines. Infection. 2016 44(1), 107–10. doi: 10.1007/s15010-015-0805-7

70. Evans DR, et al. Systematic detection of horizontal gene transfer across genera among multidrug-resistant bacteria in a single hospital. Elife 2020 doi: 10.7554/eLife.53886

71. Trobos M, Lester CH, Olsen JE, Frimodt-Møller N, Hammerum AM. Natural transfer of sulphonamide and ampicillin resistance between *Escherichia coli* residing in the human intestine. J. Antimicrob. Chemother. 2009 63, 80–86. 10.1093/jac/dkn437

72. Karami N, Martner A, Enne VI, Swerkersson S, Adlerberth I, Wold AE. Transfer of an ampicillin resistance gene between two *Escherichia coli* strains in the bowel microbiota of an infant treated with antibiotics. J Antimicrob Chemother. 2007 60(5), 1142–5.

73. Warnes SL, Highmore CJ, Keevil CW. Horizontal transfer of antibiotic resistance genes on abiotic touch surfaces: implications for public health. mBio. 2012 3(6), e00489–12. doi: 10.1128/mBio.00489-12.

74. Gibson MK, et al. Development dynamics of the preterm infant gut microbiota and antibiotic resistome. Nature Microbiology 2016 1(4), 16024. doi: 10.1038/nmicrobiol.2016.24

75. Andersson P, Engberg I, Lidin-Janson G, Lincoln K, Hull R, Hull S, Svanborg C. Persistence of *Escherichia coli* bacteriuria is not determined by bacterial adherence. Infect Immun. 1991 59(9), 2915–21.

76. Hunt M,; Mather AE, Sánchez-Busó L, Page AJ, Parkhill J, Keane JA, Harris SR. ARIBA: rapid antimicrobial resistance genotyping directly from sequencing reads. Microb Genom. 2017 3(10), e000131. doi: 10.1099/mgen.0.000131

77. Jia B, et al. CARD 2017: expansion and model-centric curation of the comprehensive antibiotic resistance database. Nucleic Acids Research. 2017 45(D1), D566–D573. doi: 10.1093/nar/gkw1004

78. Rowe WPM, Winn MD. Indexed variation graphs for efficient and accurate resistome profiling. Bioinformatics. 2018 34(21), 3601–3608. doi: 10.1093/bioinformatics/bty387

79. Phan MD, Bottomley AL, Peters KM, Harry EJ, Schembri MA. Uncovering novel susceptibility targets to enhance the efficacy of third-generation cephalosporins against ESBL-producing uropathogenic *Escherichia coli*. J Antimicrob Chemother. 2020 dkaa023 doi: 10.1093/jac/dkaa023

80. Merlin TL, Davis GE, Anderson WL, Moyzis RK, Griffith JK. Aminoglycoside uptake increased by tet gene expression. Antimicrob Agents Chemother. 1989 33(9):1549–52

81. Stavropoulos TA, Strathdee CA. Expression of the tetA(C) tetracycline efflux pump in *Escherichia coli* confers osmotic sensitivity. FEMS Microbiol. Lett. 1000 190, 147–150.

82. Heng J, Zhao Y, Liu M, Liu Y, Fan J, Wang X, Zhao Y, Zhang XC. Substrate-bound structure of the *E. coli* multidrug resistance transporter MdfA. Cell Res. 2015 25(9), 1060–73. doi: 10.1038/cr.2015.94

83. Nguyen-Distèche M, Fraipont C, Buddelmeijer N, Nanninga N. The structure and function of *Escherichia coli* penicillin-binding protein 3. Cell Mol Life Sci. 1998 54(4), 309–16.

84. Nakamura M, Maruyama IN, Soma M, Kato J, Suzuki H, Horota Y. On the process of cellular division in *Escherichia coli*: nucleotide sequence of the gene for penicillin-binding protein 3. Mol Gen Genet. 1983 191(1):1–9

85. Sun J, Deng Z, Yan A. Bacterial multidrug efflux pumps: mechanisms, physiology and pharmacological exploitations. Biochem Biophys Res Commun. 2014 453(2), 254–67. doi: 10.1016/j.bbrc.2014.05.090

86. Curtis NA, Eisenstadt RL, Turner KA, White AJ. Inhibition of penicillin-binding protein 3 of *Escherichia coli* K-12. Effects upon growth, viability and outer membrane barrier function. J Antimicrob Chemother. 1985 16(3), 287–96

87. Toh H, et al. Complete genome sequence of the wild-type commensal *Escherichia coli* strain SE15, belonging to phylogenetic group B2. Journal of Bacteriology 2010 192(4), 1165–1166.

88. Johnson JR, Porter S, Thuras P, Castanheira M. Epidemic Emergence in the United States of *Escherichia coli* Sequence Type 131-H30 (ST131-H30), 2000 to 2009. Antimicrob Agents Chemother. 2017 61(8), e00732–17. doi: 10.1128/AAC.00732-17

89. Karisik E, Ellington MJ, Pike R, Warren RE, Livermore DM, Woodford N. Molecular characterization of plasmids encoding CTX-M-15 beta-lactamases from *Escherichia coli* strains in the United Kingdom. J Antimicrob Chemother. 2006 58(3), 665–8.

90. Dudley EG, Abe C, Ghigo JM, Latour-Lambert P, Hormazabal JC, Nataro JP. An IncI1 plasmid contributes to the adherence of the atypical enteroaggregative *Escherichia coli* strain C1096 to cultured cells and abiotic surfaces. Infect Immun. 2006 74(4), 2102–14

91. Dhanji H, Doumith M, Hope R, Livermore DM, Woodford N. IS*Ecp1*-mediated transposition of linked *bla*CTX-M-3 and *bla*TEM-1b from the IncI1 plasmid pEK204 found in clinical isolates of *Escherichia coli* from Belfast, UK. J Antimicrob Chemother. 2011 66(10), 2263–5. doi: 10.1093/jac/dkr310

92. Runyen-Janecky LJ, Hong M, Payne SM. The virulence plasmid-encoded impCAB operon enhances survival and induced mutagenesis in *Shigella flexneri* after exposure to UV radiation. Infect Immun. 67(3), 1415–23.

93. Akiba M, et al. Distribution and Relationships of Antimicrobial Resistance Determinants among Extended-Spectrum-Cephalosporin-Resistant or Carbapenem-Resistant *Escherichia coli* Isolates from Rivers and Sewage Treatment Plants in India. Antimicrob. Agents Chemother. 2016 60(5), 2872–2980.

94. Carattoli A, et al. *In Silico* Detection and Typing of Plasmids using PlasmidFinder and Plasmid Multilocus Sequence Typing. 2014. Antimicrob. Agents Chemother. 2014 58(7), 3895–3903. doi: 10.1128/AAC.02412-14.

95. Purves K, Macintyre L, Brennan D, Hreggviðsson G, Kuttner E, Ásgeirsdóttir M, Young L., Green D, Edrada-Ebel R, Duncan K. Using molecular networking for microbial secondary metabolite bioprospecting. Metabolites 2016 6(1), 2.

96. Typas A, Sourjik V. Bacterial protein networks: properties and functions. Nat Rev Microbiol. 2015 13(9), 559–72. doi: 10.1038/nrmicro3508

97. Kirschner M, Gerhart J. Evolvability. Proc Natl Acad Sci USA. 1998 95(15), 8420–8427; https://doi.org/10.1073/pnas.95.15.8420

98. Szklarczyk D, et al. STRING v11: protein-protein association networks with increased coverage, supporting functional discovery in genome-wide experimental datasets. Nucleic Acids Res. 2019 47(D1), D607–D613. doi: 10.1093/nar/gky1131

99. Miryala SK, Ramaiah S. Exploring the multi-drug resistance in *Escherichia coli* O157:H7 by gene interaction network: A systems biology approach. Genomics. 2019 111(4), 958–965. doi: 10.1016/j.ygeno.2018.06.002

100. Zong Z. Complete sequence of pJIE186-2, a plasmid carrying multiple virulence factors from a sequence type 131 *Escherichia coli* O25 strain. Antimicrob Agents Chemother. 2013 57(1), 597–600. doi: 10.1128/AAC.01081-12

101. Clark G, Paszkiewicz K, Hale J, Weston V, Constantinidou C, Penn C, Achtman M, McNally A. Genomic analysis uncovers a phenotypically diverse but genetically homogeneous *Escherichia coli* ST131 clone circulating in unrelated urinary tract infections. J Antimicrob Chemother. 2012 67(4), 868–77. doi: 10.1093/jac/dkr585

102. Holden N, Totsika M, Dixon L, Catherwood K, Gally DL. Regulation of P-fimbrial phase variation frequencies in *Escherichia coli* CFT073. Infect Immun. 2007 75(7), 3325–34.

103. Spaulding CN, et al. Selective depletion of uropathogenic *E. coli* from the gut by a FimH antagonist. Nature. 2017 546(7659), 528–532. doi: 10.1038/nature22972

104. Xia Y, Gally D, Forsman-Semb K, Uhlin BE. Regulatory cross-talk between adhesin operons in *Escherichia coli*: inhibition of type 1 fimbriae expression by the PapB protein. EMBO J 2000 19, 1450–1457. https://doi.org/10.1093/emboj/19.7.1450.

105. Snyder JA, Haugen BJ, Lockatell CV, Maroncle N, Hagan EC, Johnson DE, Welch RA, Mobley HL. 2005. Coordinate expression of fimbriae in uropathogenic *Escherichia coli*. Infect Immun 73, 7588–7596. https://doi.org/10.1128/IAI.73.11.7588-7596.2005.

106. Forsman K, Göransson M, Uhlin BE. 1989. Autoregulation and multiple DNA interactions by a transcriptional regulatory protein in *E. coli* pili biogenesis. EMBO J 1989 8, 1271–1277.

107. Holden NJ, et al. Demonstration of regulatory cross-talk between P fimbriae and type 1 fimbriae in uropathogenic *Escherichia coli*. Microbiology. 2006 152(Pt4), 1143–53.

108. Brzuszkiewicz E, et al. How to become a uropathogen: comparative genomic analysis of Extraintestinal pathogenic *Escherichia coli* strains. Proc Natl Acad Sci U S A. 2006 103(34), 12879–84.

109. Welch RA, et al. Extensive mosaic structure revealed by the complete genome sequence of uropathogenic *Escherichia coli*. Proc Natl Acad Sci U S A. 2002 99(26):17020–4

110. Lillington J, Geibel S, Waksman G. Biogenesis and adhesion of type 1 and P pili. Biochim Biophys Acta. 2014 1840(9), 2783–93. doi: 10.1016/j.bbagen.2014.04.021

111. Forsberg KJ, Reyes A, Wang B, Selleck EM, Sommer MO, Dantas G. The shared antibiotic resistome of soil bacteria and human pathogens. Science. 2012 337(6098), 1107–11. doi: 10.1126/science.1220761

112. Chen SL, et al. Identification of genes subject to positive selection in uropathogenic strains of *Escherichia coli*: a comparative genomics approach. Proc Natl Acad Sci U S A. 2006 103(15), 5977–82.

113. Tenaillon O, Skurnik D, Picard B, Denamur E. The population genetics of commensal *Escherichia coli*. Nat Rev Microbiol. 2010 8(3), 207–17. doi: 10.1038/nrmicro2298

114. Didelot X, Méric G, Falush D, Darling AE. Impact of homologous and non-homologous recombination in the genomic evolution of *Escherichia coli*. BMC Genomics. 2012 13, 256. doi: 10.1186/1471-2164-13-256.

115. Dixit PD, Pang TY, Studier FW, Maslov S. Recombinant transfer in the basic genome of *Escherichia coli*. Proc Natl Acad Sci U S A. 2015 112(29), 9070–5. doi: 10.1073/pnas.1510839112

116. Tchesnokova V, Radey M, Chattopadhyay S, Larson L, Weaver JL, Kisiela D, Sokurenko EV. Pandemic fluoroquinolone resistant *Escherichia coli* clone ST1193 emerged via simultaneous homologous recombinations in 11 gene loci. Proc Natl Acad Sci U S A. 2019 116(29), 14740–14748. doi: 10.1073/pnas.1903002116

117. Smillie CS, Smith M.B, Friedman J, Cordero OX, David LA, Alm EJ. Ecology drives a global network of gene exchange connecting the human microbiome. Nature. 2011 480(7376), 241–4. doi: 10.1038/nature10571.

118. Lloyd-Price J, et al. Strains, functions and dynamics in the expanded Human Microbiome Project. Nature 2017 550(7674), 61–66.

119. Tamburini FB, Andermann TM, Tkachenko E, Senchyna F, Banaei N, Bhatt AS. Precision identification of diverse bloodstream pathogens in the gut microbiome. Nat Med. 2018 24(12), 1809–1814. doi: 10.1038/s41591-018-0202-8

120. Sundén F, Håkansson L, Ljunggren E, Wullt B. *Escherichia coli* 83972 bacteriuria protects against recurrent lower urinary tract infections in patients with incomplete bladder emptying. J Urol. 2010 184(1), 179–85. doi: 10.1016/j.juro.2010.03.024

121. Sundén F, Håkansson L, Ljunggren E, Wullt B. Bacterial interference—is deliberate colonization with *Escherichia coli* 83972 an alternative treatment for patients with recurrent urinary tract infection? Int J Antimicrob Agents. 2006 28 Suppl 1, S26–9.

122. Watts RE, et al. *Escherichia coli* 83972 expressing a P fimbriae oligosaccharide receptor mimic impairs adhesion of uropathogenic *E. coli*. J Infect Dis. 2012 206(8), 1242–9.

123. Zdziarski J, et al. Host imprints on bacterial genomes--rapid, divergent evolution in individual patients. PLoS Pathog. 2010 6(8), e1001078. doi: 10.1371/journal.ppat.1001078

124. Duprilot M, et al. Success of *Escherichia coli* O25b:H4 ST131 clade C associated with a decrease in virulence. BioRxiv 2019 doi:https://doi.org/10.1101/786350.

125. Mora A, et al. Virulence patterns in a murine sepsis model of ST131 *Escherichia coli* clinical isolates belonging to serotypes O25b:H4 and O16:H5 are associated to specific virotypes. PLoS ONE 2014 9, e87025.

126. Fondi M, Karkman A, Tamminen MV, Bosi E, Virta M, Fani R, Alm E, McInerney JO. Every Gene Is Everywhere but the Environment Selects: Global Geolocalization of Gene Sharing in Environmental Samples through Network Analysis. Genome Biol Evol. 2016 8(5), 1388–400. doi: 10.1093/gbe/evw077

127. Downing T. Tackling Drug Resistant Infection Outbreaks of Global Pandemic *Escherichia coli* ST131 Using Evolutionary and Epidemiological Genomics. Microorganisms. 2015 3(2), 236–67. doi: 10.3390/microorganisms3020236

128. Leggett RM, et al. Rapid MinION profiling of preterm microbiota and antimicrobial-resistant pathogens. Nat Microbiol 2020 5, 430–442 https://doi.org/10.1038/s41564-019-0626-z.

129. Smith H, et al. Characterization of epidemic IncI1-Iγ plasmids harboring Ambler class A and C genes in *Escherichia coli* and *Salmonella enterica* from animals and humans. Antimicrob Agents Chemother 2015 59, 5357–5365. https://doi.org/10.1128/AAC.05006-14.

130. Rozwandowicz M, Brouwer MSM, Fischer J, Wagenaar JA, Gonzalez-Zorn B, Guerra B, Mevius DJ, Hordijk J. 2018. Plasmids carrying antimicrobial resistance genes in Enterobacteriaceae. J Antimicrob Chemother 73:1121–1137. https://doi.org/10.1093/jac/dkx488

131. Rodríguez-Navarro J, Miró E, Brown-Jaque M, Hurtado JC, Moreno A, Muniesa M, González-López JJ, Vila J, Espinal P, Navarro F. Comparison of Commensal and Clinical Isolates for Diversity of Plasmids in *Escherichia coli* and *Klebsiella pneumoniae*. Antimicrob Agents Chemother. 2020 64(5). pii: e02064–19. doi: 10.1128/AAC.02064-19

132. Dimitriu T, Medaney F, Amanatidou E, Forsyth J, Ellis RJ, Raymond B. Negative frequency dependent selection on plasmid carriage and low fitness costs maintain extended spectrum β-lactamases in *Escherichia coli*. Sci Rep. 2019 9(1), 17211. doi: 10.1038/s41598-019-53575-7

133. Totsika M, et al. Insights into a multidrug resistant *Escherichia coli* pathogen of the globally disseminated ST131 lineage: genome analysis and virulence mechanisms. PLoS One 2011 6, e26578.

134. Schembri MA, Zakour NL, Phan MD, Forde BM, Stanton-Cook M, Beatson SA. Molecular Characterization of the Multidrug Resistant *Escherichia coli* ST131 Clone. Pathogens. 2015 4(3), 422–30. doi: 10.3390/pathogens4030422

135. Wurpel DJ, et al. Comparative proteomics of uropathogenic *Escherichia coli* during growth in human urine identify UCA-like (UCL) fimbriae as an adherence factor involved in biofilm formation and binding to uroepithelial cells. J. Proteomics 2016 131, 177–189.

136. Lane MC, Mobley HL. Role of P-fimbrial-mediated adherence in pyelonephritis and persistence of uropathogenic *Escherichia coli* (UPEC) in the mammalian kidney. Kidney Int 2007 72, 19–25. https://doi.org/10.1038/sj.ki.5002230.

137. Gunther NW, Lockatell V, Johnson DE, Mobley HLT. *In vivo* dynamics of type, 1 fimbria regulation in uropathogenic *Escherichia coli* during experimental urinary tract infection. Infect Immun 2001 69, 2838–2846.

138. Holden NJ, Uhlin BE, Gally D.. PapB paralogues and their effect on the phase variation of type 1 fimbriae in *Escherichia coli*. Mol Microbiol. 2001 42(2), 319–30

139. Holden NJ, Gally D. Switches, cross-talk and memory in *Escherichia coli* adherence. J Med Microbiol. 2004 53(Pt 7), 585–593. doi: 10.1099/jmm.0.05491-0

140. Whitmer GR, Moorthy G, Arshad M. The pandemic *Escherichia coli* sequence type 131 strain is acquired even in the absence of antibiotic exposure. PLoS Pathog. 2019 15(12), e1008162. doi: 10.1371/journal.ppat.1008162

141. NIH HMP Working Group, et al. The NIH Human Microbiome Project. Genome Res. 2009 19(12), 2317–23. doi: 10.1101/gr.096651.109.

142. Rudick CN, Taylor AK, Yaggie RE, Schaeffer AJ, Klumpp DJ. Asymptomatic Bacteriuria *Escherichia coli* are Live Biotherapeutics for UTI. PLoS One 2014 9(11), 1–9.

143. Hancock V, Seshasayee AS, Ussery DW, Luscombe NM, Klemm P. Transcriptomics and adaptive genomics of the asymptomatic bacteriuria Escherichia *coli* strain 83972. Mol Genet Genomics. 2008 279(5), 523–34. doi: 10.1007/s00438-008-0330-9.

144. Floyd RV, Upton M, Hultgren SJ, Wray S, Burdyga TV, Winstanley C. *Escherichia coli*-mediated impairment of ureteric contractility is uropathogenic *E. coli* specific. J. Infect. Dis. 2012, 206, 1589–1596.

145. Bolger AM, Lohse M, Usadel B. Trimmomatic: a flexible trimmer for Illumina sequence data. Bioinformatics. 2014 30(15), 2114–20. doi: 10.1093/bioinformatics/btu170

146. Li H, et al. The Sequence Alignment/Map format and SAMtools. Bioinformatics. 2009 25(16), 2078–9. doi: 10.1093/bioinformatics/btp352

147. Jones P, et al. InterProScan 5: genome–scale protein function classification. Bioinformatics. 2014 30(9), 1236–40. doi: 10.1093/bioinformatics/btu031

148. Carver T, Berriman M, Tivey A, Patel C, Böhme U, Barrell BG, Parkhill J, Rajandream MA. Artemis and ACT: viewing, annotating and comparing sequences stored in a relational database. Bioinformatics. 2008 24(23), 2672–6. doi: 10.1093/bioinformatics/btn529.

149. Sievers F, et al. Fast, scalable generation of high-quality protein multiple sequence alignments using Clustal Omega. Molecular Systems Biology 2011 7, 539 doi: 10.1038/msb.2011.75

150. Nicolau M. Levine AJ, Carlsson G. Topology based data analysis identifies a subgroup of breast cancers with a unique mutational profile and excellent survival. Proc Natl Acad Sci U S A. 2011 108(17), 7265–70. doi: 10.1073/pnas.1102826108

151. Vietoris L. Über den höheren Zusammenhang kompakter Räume und eine Klasse von zusammenhangstreuen Abbildungen. Mathematische Annalen 1927 97(1), 454–472.

152. Rahm A. HomologyLive. 2019 http://math.uni.lu/~rahm/HomologyLive.html.

153. LinBox, The LinBox Group. Exact Linear Algebra over the Integers and Finite Rings, Version 1.1.6, 2008.

154. Kim SR, Komano T. The plasmid R64 thin pilus identified as a type IV pilus. J Bacteriol. 1997 179(11), 3594–603.

