## Supplementary material for "Plasmids shape the diverse accessory resistomes of *Escherichia coli* ST131": Suppl_Data

### Supplementary Data


| Page | Item | Title |
|---|---|---|
| A shared core and diverse accessory preterm infant resistome in *E. coli* | | |
| 1 | **Table S1** | The 16 antibiotics used to identify 794 AMR genes on contigs originally from preterm infants. |
| 1 | **Table S2** | The five AMR genes unique to all ST131 Clade C genomes except 8289_1#27 encoded *bla*$_{TEM}$ genes (an Ambler class A β-lactamase). |
| Extensive plasmid rearrangements in closely related ST131 Clade C genomes | | |
| 2 | **Table S3** | PlasmidFinder Inc group alignments against pV130 and seven ST131 isolates. |
| 3 | **Table S4** | Key known AMR, plasmid persistence and conjugation genes for pEK204, pEK499, pEK516, pCA14, pV130a and pV130b. |
| 4 | **Figure S1** | Comparison of pEK516 (64,471 bp) with ST131 and 83972. |
| 5-6 | **Figure S2** | Read mapping distributions for seven ST131 to pV130 contigs. |
| 7 | **Figure S3** | Comparison of plasmid pCA14 (155 Kb) with seven ST131 C1/C2 isolates. |
| Higher rates of pEK499, pEK516 and pEK204 protein interactions with chromosomal proteins | | |
| 8 | **Table S5** | The PPIN characteristics of the protein-coding genes from Miryala et al (2018). |
| 8 | **Table S6** | The seven plasmids' accessions, numbers of genes with interactions given a combined score of 400+, Inc groups, conjugative ability and AMR genes. |
| 9 | **Figure S4** | The ratio of non-trivial loops per PPI (y-axis) versus the STRING combined score (x-axis) for all 60 genes from Miryala et al (2018) Table 1 [99]. |
| 9 | **Figure S5** | The ratio of non-trivial loops per interaction versus the STRING combined score for all chromosomal genes, all genes on pEK499 and all those on pEK204. |
| 9 | **Figure S6** | The ratio of interactions per protein for genes encoded on the chromosome or the pEK499 and pEK204 plasmids across the STRING combined score. |
| ST131 genomes had an ancestral pEK499-like plasmid but some gained a pEK204-like one | | |
| 11 | **Figure S7** | The distributions of pEK204-like and pEK499-like regions in ST131 assemblies. |
| 11 | **Figure S8** | The distributions of pEK204-like regions in ST131 Clade C genome assemblies. |
| 12 | **Table S7** | Read library summary statistics for each ST131 sample and reference genome. |
| 12 | **References** | (not in main text) |

### A core preterm infant resistome within across E. coli isolates

| Antibiotic | Class | Abbreviation | MIC (ug/ml) |
|---|---|---|---|
| Amoxicillin | β-lactam | AX | 16 |
| Amoxicillin+Clavulanate | β-lactam | AXCL | 16-8 |
| Ampicillin | β-lactam | AP | 64 |
| Aztreonam | β-lactam | AZ | 8 |
| Cefepime | β-lactam | CP | 8 |
| Cefoxitin | β-lactam | CX | 64 |
| Ceftazidime | β-lactam | CZ | 16 |
| Chloramphenicol | amphenicol | CH | 8 |
| Ciprofloxacin | quinolone | CI | 0.5 |
| Colistin | polymixin | COL | 8 |
| Gentamicin | aminoglycoside | GE | 16 |
| Meropenem | Carbapenem | ME | 16 |
| Penicillin-G | β-lactam | PE | 128 |
| Piperacillin | β-lactam | PI | 16 |
| Tetracycline | tetracycline | TE | 8 |
| Tigecycline | tetracycline | TG | 2 |

**Table S1**. The 16 antibiotics used to identify 794 AMR genes from functional metagenomic screening [74]. This reference resistome came from 21 infants' faecal samples that were assembled together as 2,004 redundant contigs following experimentally testing for resistance to these 16 antibiotics and Illumina HiSeq sequencing [74]. This approach sheared metagenomic DNA into fragments ligated in *E. coli* that were grown on media with these antibiotic concentrations [74]. A total of 336 (16x21) transformants were attempted, of which 183 (54%) were successful [74]. From the colonies that grew enough during 18 hours, they found 2,004 contigs (longer than 0.5 Kb) with 794 unique genes [74].

### A shared core and diverse accessory preterm infant resistome in *E. coli*

| Contig ID (length) | Gene (gene product function) and location on contig | #Contigs with gene | Contig antibiotic profile |
|---|---|---|---|
| 1_C6_AXC.3 (1,647 bp) | *APH(3'')-Ib* (aminoglycoside phosphotransferase) at bases 2-811 * | 4 | Amoxicillin + Clavulanate |
| | *APH(6')-Id* (aminoglycoside phosphotransferase) at bases 811-1,647 | 2 | |
| 1_C6_AP.7 (7,018 bp) | *APH(3'')-Ib* (aminoglycoside phosphotransferase) at bases 5,637-6,440 * | 4 | Ampicillin |
| | *APH(6')-Id* (aminoglycoside phosphotransferase) at bases 4,801-5,637 | 2 | |
| 5_D1_AXCL. 8 (9,168 bp) | *tetA* (tetracycline efflux pump) at bases 2,805-3,221 | 1 | Amoxicillin + Clavulanate |
| | *tetD* (tetracycline efflux pump) at bases 788-2,011 ** | 2 | |

**Table S2.** Four AMR genes unique to 8289_1#24 were derived from six contigs, three of which are listed here. The other three were: * a *APH(3'')-Ib* gene was at bases 166-969 on a 6,254 bp-long contig 1_C6_AX.4 associated with amoxicillin-resistance, and at bases 441-1,244 on a 2,077 bp-long contig 1_C6_PE.1 associated with penicillin-G-resistance; and ** a *tetD* gene was at bases 1,850-2,266 on a 3,300 bp-long contig 5_D1_AP.6 associated with ampicillin-resistance.

| **Replicon ID** | **8289_1 #27** | **8289_1 #60** | **8289_1 #24** | **8289_1 #3** | **8289_1 #34** | **8289_1 #35** | **8289_1 #91** | **pV130a contigs** | **pV130b contigs** |
|---|---|---|---|---|---|---|---|---|---|
| **Subclade** | **C2_9** | **C2_9** | **C2_8** | **C2_7** | **C2_7** | **C1** | **C1** | | |
| IncFI | 1 | 1 | 1 | | 1 | 1 | 1 | 1 | 1*** |
| IncFIA | 1 | 1 | 1 | 1 | 1 | 1 | 1 | 1 | |
| IncFIB | 1 | 1 | 1 | | | 1 | 1 | 1** | |
| IncFII | | | | | | 1 | 1 | | |
| IncX1 | | | | | | 1 | 1 | | |
| IncQ1 | | | 1 | | 1 | 1 | | | |
| IncI1 | 1 | | | | | | | | |
| IncX4 | | 1 | | | | | 1 | | |
| IncX3 | | | | | | | | | 1 |
| Col156 | 1 | 1 | | | | 1 | 1 | 1* | |
| Col_BS51 | 1 | 1 | 1 | | 1 | 1 | | | |
| Col_MG8 | 1 | 1 | 1 | 1 | 1 | 1 | 1 | | |
| ColRNAI | | | | 1 | | | | | |

**Table S3**. PlasmidFinder Inc group assignments for pV130 and seven ST131 isolates. Values indicate presence of the replicon type or element. Matches for pV130a shown for scaffold a_1, except for * in a_4 and ** in a_2. Matches for pV130a shown for scaffold a_1, except *** in b_3. The IncF1B replicon was IncFIB_AP00918.

| **AMR genes** | **Description** | **pEK204** | **pEK499** | **pEK516** | **pCA14** | **pV130a** | **pV130b** |
|---|---|---|---|---|---|---|---|
| *aac(3)-II* | aminoglycoside N(3')-acetyltrans-ferase III; gentamicin-R, netilmicin-R, tobramycin-R, sisomicin-R | | | 1 | | | |
| *aac(6')-Ib-cr* | aminoglycoside N(6')-acetyltrans-ferase type Ib-cr; quinolone-R | | 1 | 1 | 1 | | 1 |
| *aadA5* | aminoglycoside resistance protein | | 1 | | 1 | | |
| *catB4* | chloramphenicol acetyltransferase; inactivates chloramphenicol | | 1 | 1 | | | |
| *CTX-M* | extended spectrum beta-lactamase | 1 | 1 | 1 | 1 | 1 | |
| *dfrA7* | dihydrofolate reductase type VII; trimethoprim resistance | | 1 | | | | |
| *mph(A)* | macrolide 2-phosphotransferase; inactivates erythromycin | | 1 | | 1 | 1 | 1 |
| *OXA-1* | extended spectrum beta-lactamase | | 1 | 1 | 1 | | 1 |
| *sul1* | dihydropteroate synthase; sulfonamide resistance protein | | 1 | | 1 | | |
| *TEM-1* | beta-lactamase | 1 | 1 | 1 | | | |
| *tetA* | tetracycline resistance protein | | 1 | 1 | 1 | | |
| ***Persistence genes*** | **Description** | **pEK204** | **pEK499** | **pEK516** | **pCA14** | **pV130a** | **pV130b** |
| *ccdA* | plasmid maintenance protein | | 1 | | 1 | 1 | |
| *ccdB* | plasmid maintenance protein | | 1 | | 1 | 1 | |
| *hok* | post-segregation killing protein | | 1 | 1 | 1 | | 1 |
| *mok* | modulator of Hok | | 1 | 1 | 1 | | 1 |
| *parM* | plasmid segregation protein | 1 | | 1 | | | 1 |
| *pemI* | stable plasmid inheritance transcriptional regulator/antitoxin | | 1 | 1 | 1 | 1 | 1 |
| *pemK* | stable plasmid inheritance protein toxin-antitoxin system toxin | | 1 | 1 | 1 | 1 | 1 |
| *stbB* | stable plasmid inheritance protein B | 1 | | 1 | | | 1 |
| *vagC* | virulence-associated protein vagC; toxin addiction system; antitoxin | | 1 | | 1 | 1 | |
| *vagD* | virulence-associated protein vagD; toxin addiction system; antitoxin | | 1 | | 1 | 1 | |
| ***Conjugation genes*** | **Description** | **pEK204** | **pEK499** | **pEK516** | **pCA14** | **pV130a** | **pV130b** |
| *traC* | conjugal transfer ATP-binding protein; conjugal transfer | 1 | 1 | | 1 | | |
| *traX* | responsible for N-terminal acetyl-ation of F pilin; F pilus assembly | 1 | | | 1 | | |

**Table S4.** Key known AMR, plasmid persistence and conjugation genes for pEK204, pEK499, pEK516, pCA14, pV130a and pV130b based on alignment with BLAST to CARD with confirmation using the annotation files. 1 indicates presence. PEK499 is stably inherited because it has post-segregation killing genes *hok* and modulator *mok*, toxin-antitoxin system genes (*pemI-pemK*, *ccdA-ccdB*), and two copies of virulence-associated genes, *vagC* and *vagD*. $Bla_{\mathrm{TEM\text{-}1b}}$ was 860 bp in pEK204 and 728 bp in pEK499 and pEK516. *ParM* and *StbB* were 980 bp in pEK204 but 962 bp in pEK499 and pEK516.

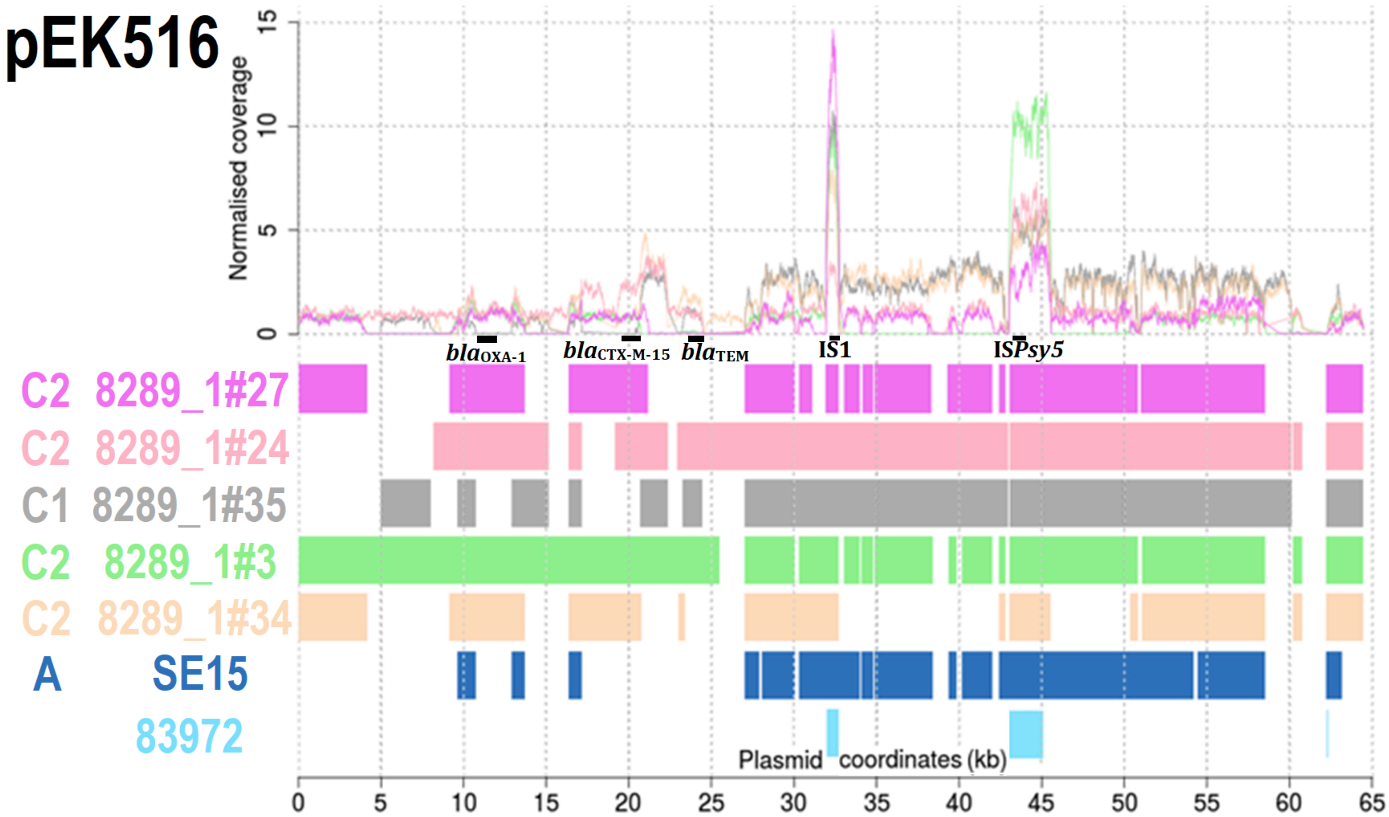


**Figure S1**. Comparison of pEK516 (64,471 bp) with five ST131 Subclade C1 and C2 isolates, Clade A reference SE15 (navy) and 83972 (cyan). Top: normalised coverage of mapped reads for the five ST131 Clade C isolates showed high copy numbers at IS*1* (32 Kb) and IS*Psy5* (with unannotated genes at 43-45 Kb) (shown by black boxes). Bottom: BLAST alignment similarity showed limited showed limited matching for 83972 and SE15, relative to the Subclade C1 and C2 assemblies: 8289_1#27 from C2_9 (mauve), 8289_1#24 from C2_8 (pink), 8289_1#35 from C1 (grey), 8289_1#3 from C2_7 (light green) and 8289_1#34 from C2_7 (beige). Genes encoding $bla_{\text{OXA-1}}$, $bla_{\text{CTX-M-15}}$ and $bla_{\text{TEM}}$ were at 12, 20 and 24 Kb (respectively) highlighted with black boxes. Plasmid pEK516 lacks *traX* and *traC*, but has *traA/B/D/E/K/L/M/P/V/Y*. Matches spanning >300 bp are shown. The region at 22-61 Kb that is inverted relative to pEK499 was largely present in the ST131 here. 8289_1#34 had limited homology at the *tra* region (33-40 Kb).

**Figure S2** (below and overleaf). Read mapping distributions for seven ST131 to pV130 contigs showing (top of each panel) the normalised read coverage and (bottom) the presence of regions as coloured bars for 8289_1#27 from C2_9 (pink), 8289_1#60 from C2_9 (mauve), 8289_1#24 from C2_8 (light pink), 8289_1#35 from C1 (grey), 8289_1#91 from C1 (blue), 8289_1#3 from C2_7 (green) and 8289_1#34 from C2_7 (biege) for (A) pV130a_1 with a *vgaC* gene (encoding a ABC-F subfamily protein associated with streptogramin-resistance) absent in C1 and IncF2A replicon present in all; (B) pV130a_5 with two IS26 copies, a *mph(A)* gene (encoding a macrolide 2-phosphotrans-ferase that inactivates erythromycin) and a *Mrx* gene (encoding a macrolide phosphotransferase assisting MphA); (C) pV130b_1 with an IS26; (D) pV130b_3 with an IS26 and an IncF2 replicon; (E) pV130b_23 with two IS26 copies; (F) pV130a_2 with an IS26 and IncF1B replicon present in all; (G) pV130a_7 with two IS26 copies and a $bla_{CTX\text{-}M\text{-}15}$ gene (position 837-1,713) in C2 only; (H) pV130b_2 with two IS26 copies, a bleomycin resistance protein (BRP) metallo-beta-lactamase (MBL) gene in C1 and C2-8 only, and a $bla_{\mathrm{NDM}}$ gene (position 13,589-14,146) in C1 only; (I) pV130a_22 with two IS26 copies, a *mph(A)* gene and a Mrx gene; and (J) pV130b_190 with two IS26 copies and a region in C2 only with a *catB3* gene (encoding a chloramphenicol acetyltransferase that inactivates phenicols), an $bla_{\mathrm{OXA\text{-}1}}$ gene and quinolone-resistance gene *aac(6')-Ib-cr*. Plasmid pV130 was isolated by [93] from sewage treatment plant water in India that found 49 plasmids of which two were pV130a and pV130b, with a total of 15 contigs. The former was 108,055 bp, has nine contigs, was $bla_{\mathrm{CTX\text{-}M\text{-}15}}$-positive, contained 134 genes and had replicon types FIA, FIB and FII. Plasmid pV130b was 78,386 bp, had six scaffolds, was $bla_{\mathrm{NDM}}$-positive, had 111 genes and replicon type FII. Both have six plasmid persistence genes each and lack *traC* and *traX* .

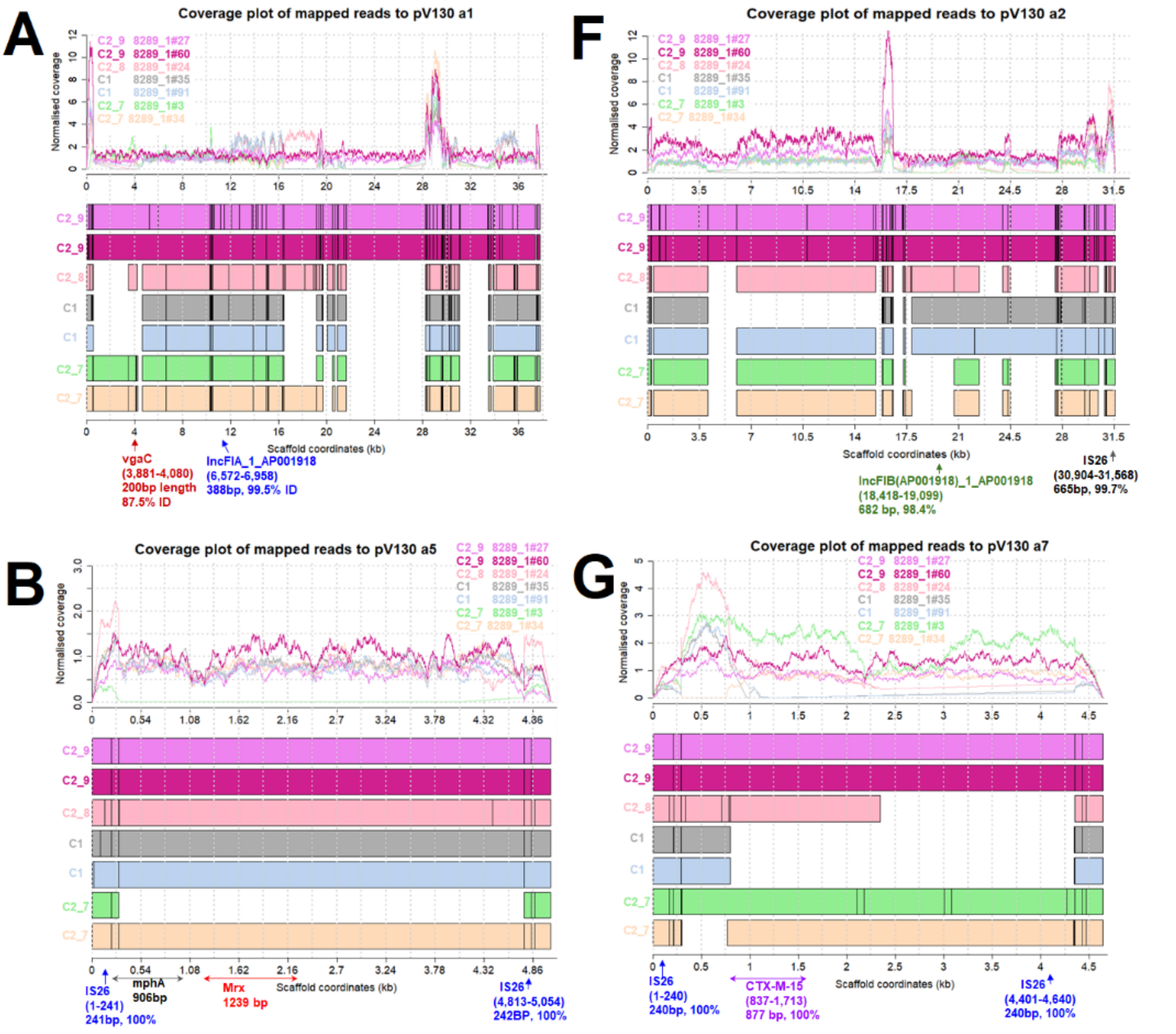

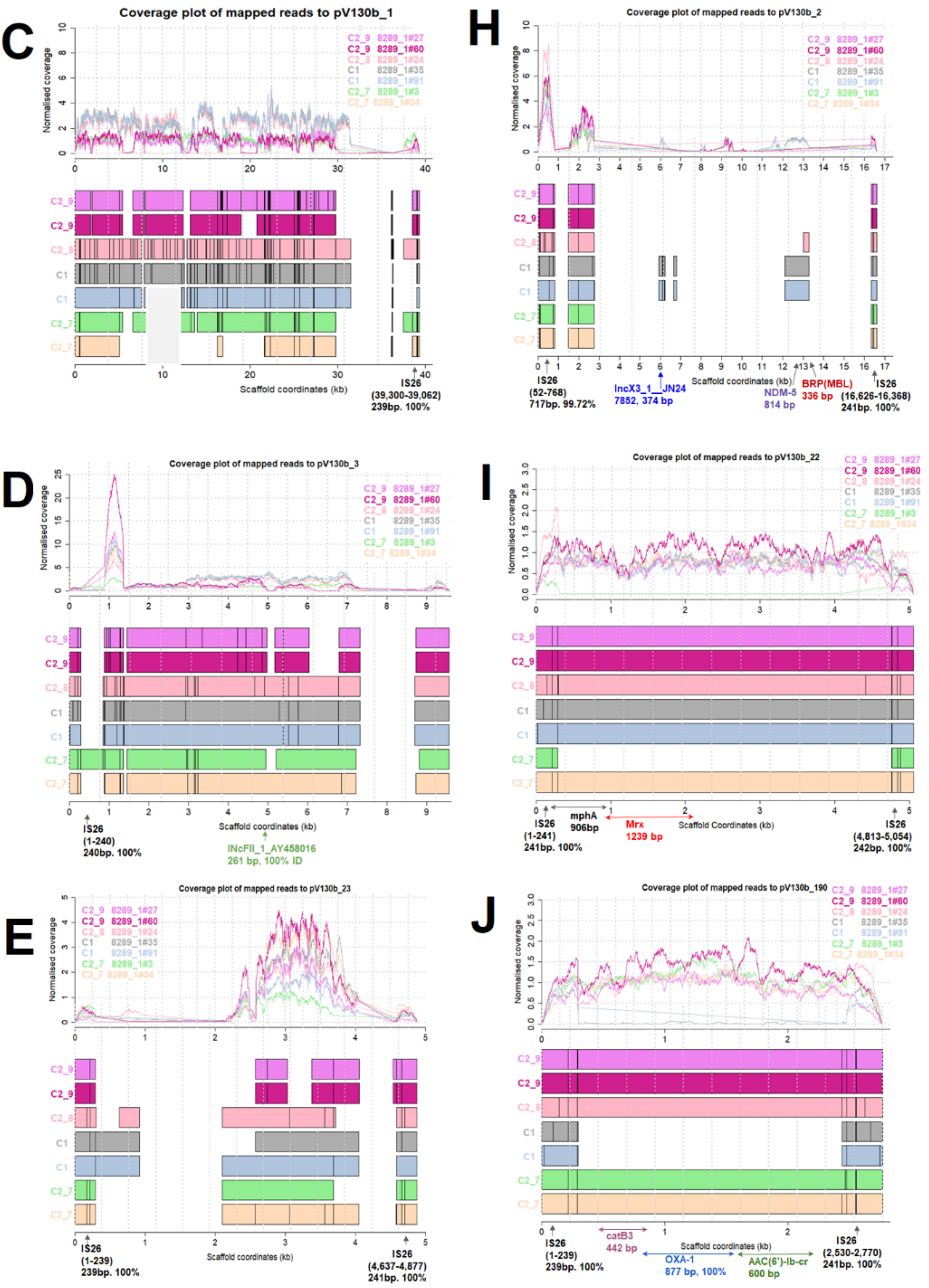
C
Coverage plot of mapped reads to pV130b_1
Normalised coverage
C2_9 8289_1#27
C2_9 8289_1#60
C2_8 8289_1#24
C1 8289_1#35
C1 8289_1#91
C2_7 8289_1#3
C2_7 8289_1#34
Scaffold coordinates (kb)
IS26
(39,300-39,062)
239bp. 100%
H
Coverage plot of mapped reads to pV130b_2
IS26
(52-768)
717bp. 99.72%
IncX3_1__JN24
7852, 374 bp
Scaffold coordinates (kb)
NDM-5
814 bp
BRP(MBL)
336 bp
IS26
(16,626-16,368)
241bp. 100%
D
Coverage plot of mapped reads to pV130b_3
IS26
(1-240)
240bp. 100%
Scaffold coordinates (kb)
IncFII_1_AY458016
261 bp, 100% ID
I
Coverage plot of mapped reads to pV130b_22
IS26
(1-241)
241bp. 100%
mphA
906bp
Mrx
1239 bp
Scaffold Coordinates (kb)
IS26
(4,813-5,054)
242bp. 100%
E
Coverage plot of mapped reads to pV130b_23
IS26
(1-239)
239bp. 100%
Scaffold coordinates (kb)
IS26
(4,637-4,877)
241bp. 100%
J
Coverage plot of mapped reads to pV130b_190
IS26
(1-239)
239bp. 100%
catB3
442 bp
OXA-1
877 bp, 100%
Scaffold coordinates (kb)
AAC(6')-Ib-cr
600 bp
IS26
(2,530-2,770)
241bp. 100%

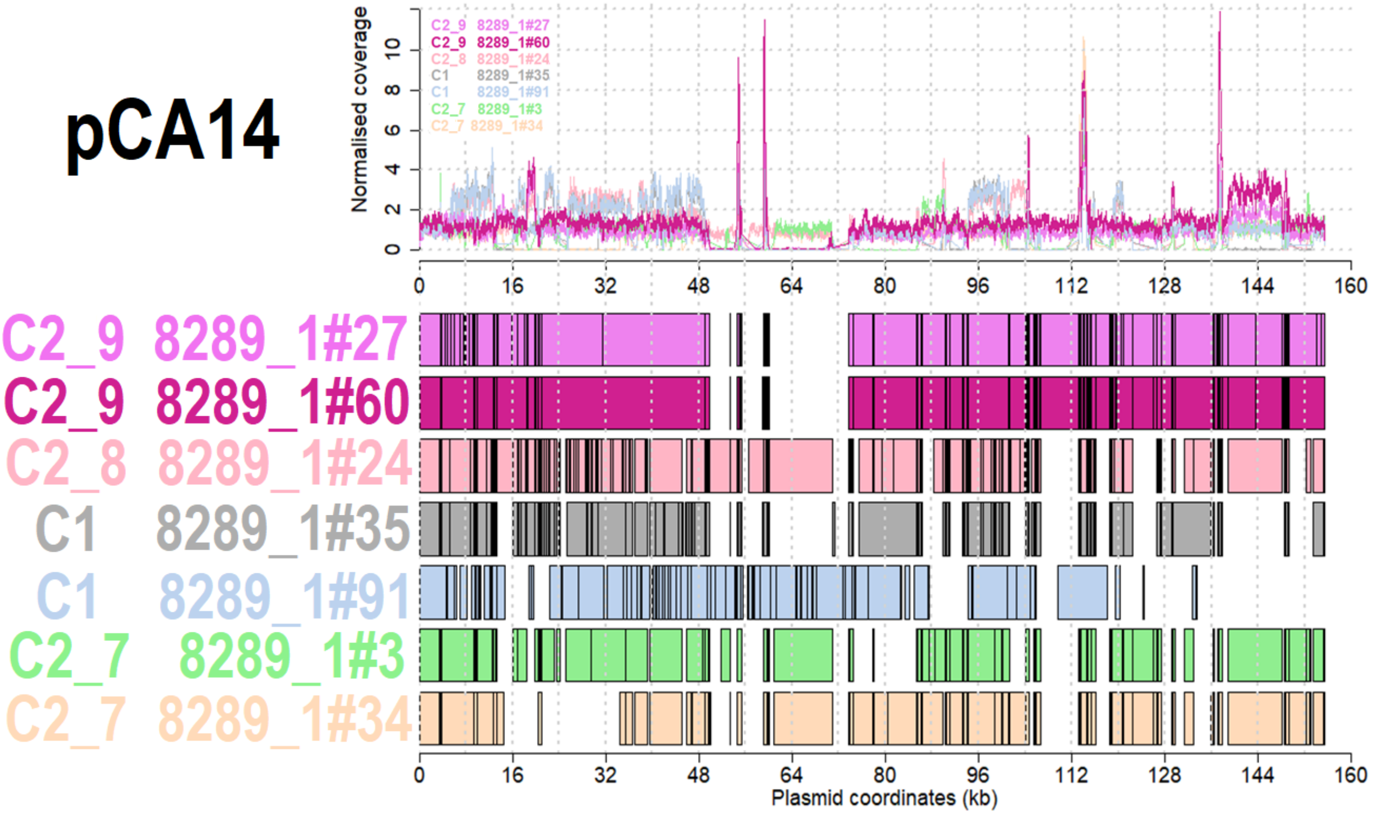


**Figure S3**. Comparison of F31:A4:B1 plasmid pCA14 (155 Kb) with seven ST131 Clade C1 and C2 isolates. Reads for all seven ST131 were mapped to conjugative plasmid pCA14, is structurally similar to pEK499, has seven known AMR genes and eight plasmid persistence genes [59] (Table S9). Top: normalised coverage of mapped reads for the seven read libraries a showed high copy number at different regions. Bottom: Regions of similarity based on BLAST alignments showed variable matching for the samples: 8289_1#27 from C2_9 (pink), 8289_1#60 from C2_9 (mauve), 8289_1#24 from C2_8 (light pink), 8289_1#35 from C1 (grey), 8289_1#91 from C1 (blue), 8289_1#3 from C2_7 (green) and 8289_1#34 from C2_7 (beige). All seven libraries had extensive similarity but a mosaic structure, and all bar 8289_1#3 possessed the *mph(A)* and *Mrx* genes (associated with erythromycin resistance).

**Higher rates of pEK499, pEK516 and pEK204 protein interactions with chromosomal proteins**

| Subset name | #genes | #PPIs within group | #PPIs with chromosome | #PPIs per protein (within group) | #PPIs per protein (chromosome) |
|---|---|---|---|---|---|
| All | 60 | 229 | 2,307 | 3.8 | 10.1 |
| Cluster_1 | 30 | 138 | 1,125 | 4.6 | 8.2 |
| All_AMR | 19 | 57 | 852 | 4.0 | 14.9 |

**Table S5**. The PPI network connectivity of genes from Table 1 of [99]. All were unique and had PPI data (bar six in the full set of 60). The numbers of PPIs within each cluster is shown, followed by the numbers per group with the chromosome. The TDA-based approach was tested using results from a previous clustering analysis that had five clusters with 60 proteins in total (see Table 1 of [99]). We examined the full set of 60 (Figure S8), cluster 1 alone with 30 proteins to examine intra-cluster patterns, and 19 AMR-related proteins from all five clusters to look at inter-cluster trends. This showed that the within-cluster interactions per protein (3.8 to 4.6) and chromosomal interactions per protein (8.2 to 14.9) were consistent across groups.

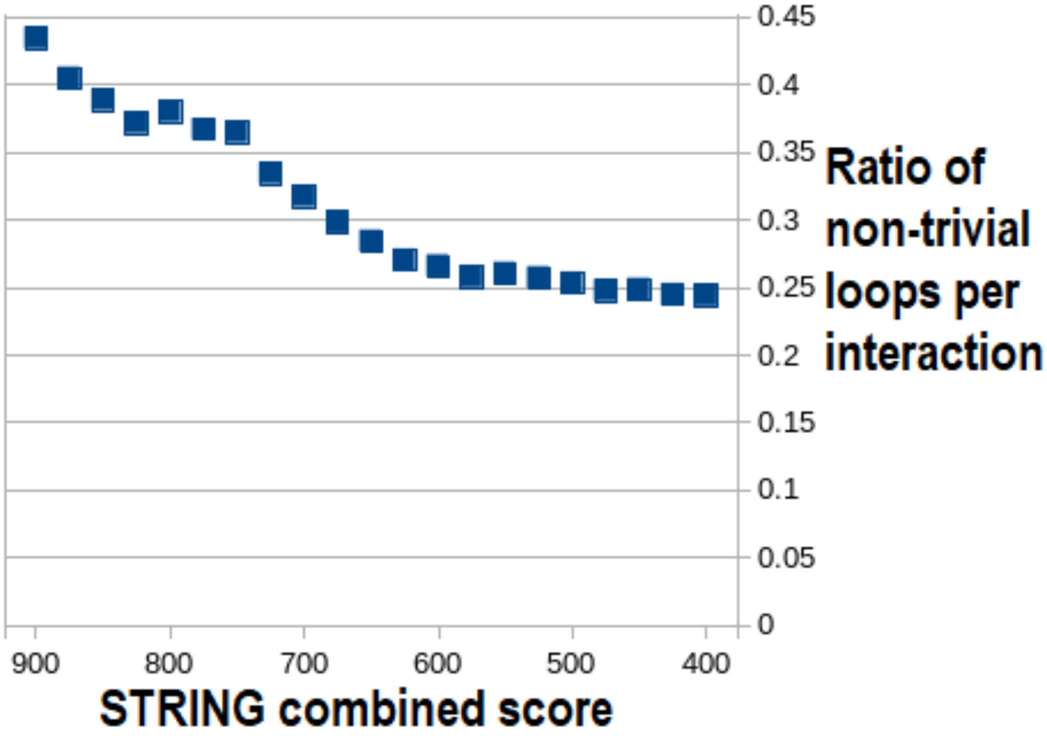


**Figure S4.** The ratio of non-trivial loops per PPI (y-axis) versus the STRING combined score (x-axis) for all 60 genes from Table 1 of [99]. This showed that the non-trivial loops per PPI (missing interactions) increased with the combined score.

| Plasmid | Accession | #genes with PPIs | Inc group(s) | Conjugative (intact *tra*) | AMR |
|---|---|---|---|---|---|
| pEK204 | EU935740 | 24 | I1 | Yes | Yes |
| pEK499 | EU935739 | 26 | F2/F1A | No | Yes |
| pEK516 | EU935738 | 26 | F2A | No | Yes |
| pJIE186-2 | NC 020271 | 7 | F1B/F2A/F1A | No | No |
| pCA14 | CP009231 | 14 | F2/F1A/F1B | Yes | Yes |
| pEC958A | HG941719 | 13 | F1A/F2 | No | Yes |
| pECSF1 | NC 013655 | 3 | F2A/FIB | Yes | No |

**Table S6.** The seven plasmids' accessions, numbers of genes with interactions given a combined score of >400, Inc groups, conjugative ability and presence of AMR genes. PJIE186-2's IncF1A was partial.

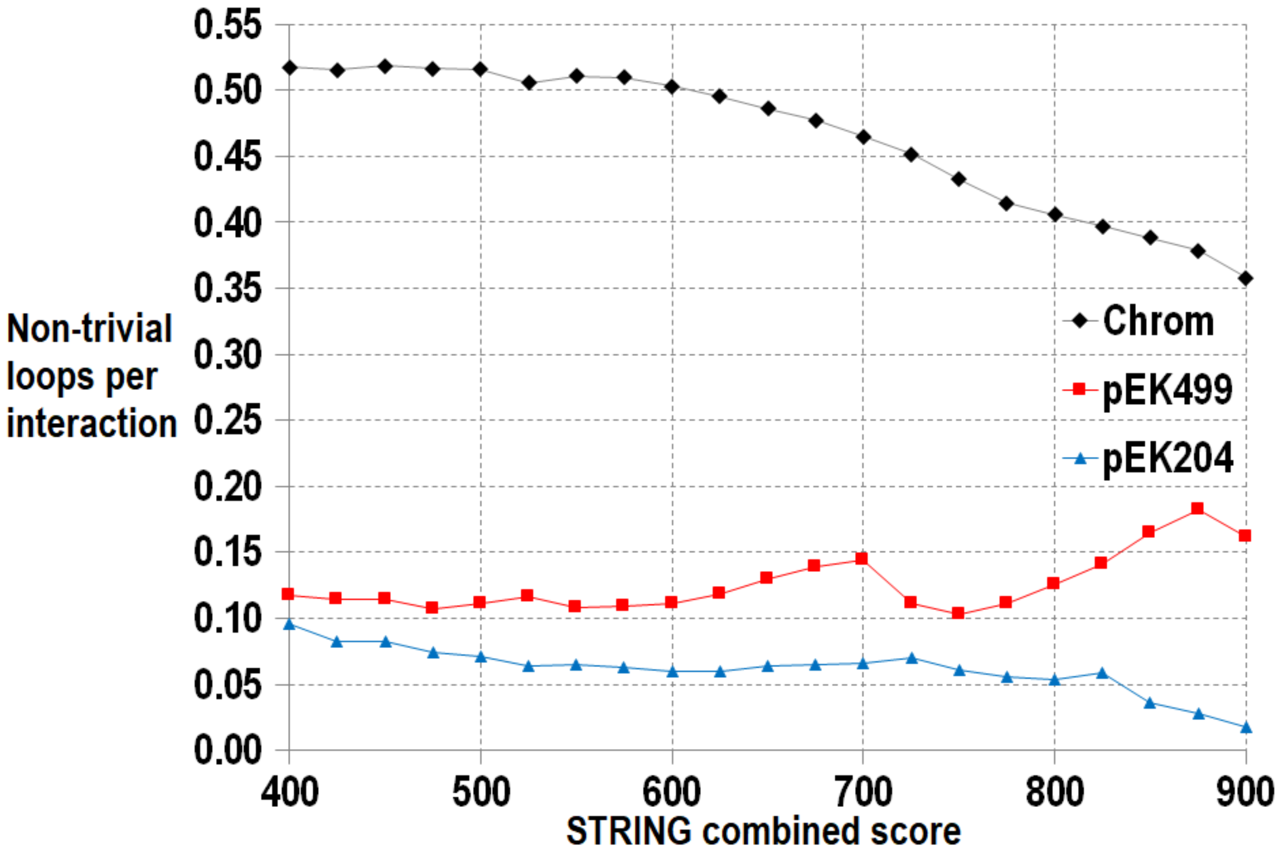


**Figure S5**. The ratio of non-trivial loops per PPI (y-axis) versus the STRING combined score (x-axis) for all *E. coli* 4,146 chromosomal genes (black), all genes on pEK499 (red) and all those on pEK204 (blue). This showed that the non-trivial loops per PPI (missing interactions) varied little for these plasmids with respect to the STRING combined score.

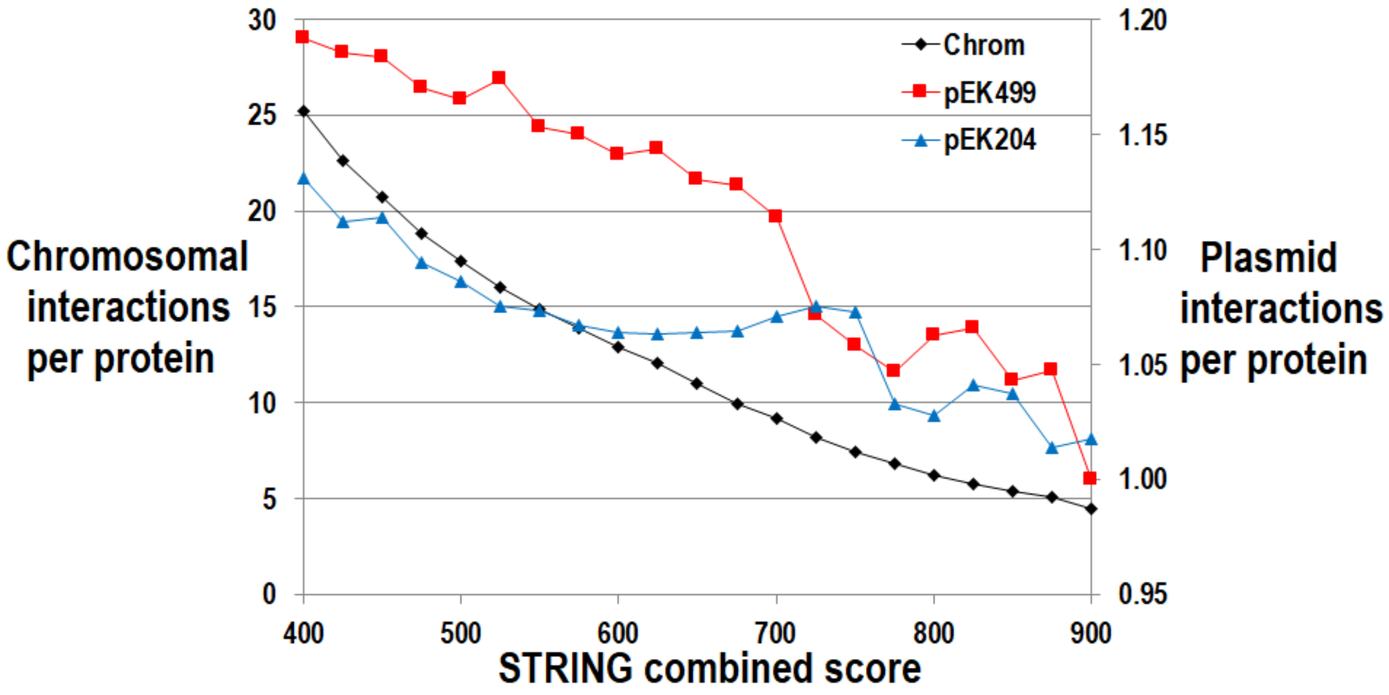


**Figure S6.** The ratio of PPIs per protein for genes encoded on the chromosome in black (left y-axis) or the pEK499 in red and pEK204 in blue plasmids (right y-axis, note difference in scale) versus the STRING combined score (x-axis). As the combined score threshold became higher, the decay of PPIs per protein was about equivalent across datasets, showing the STRING combined score was not a confounder here.

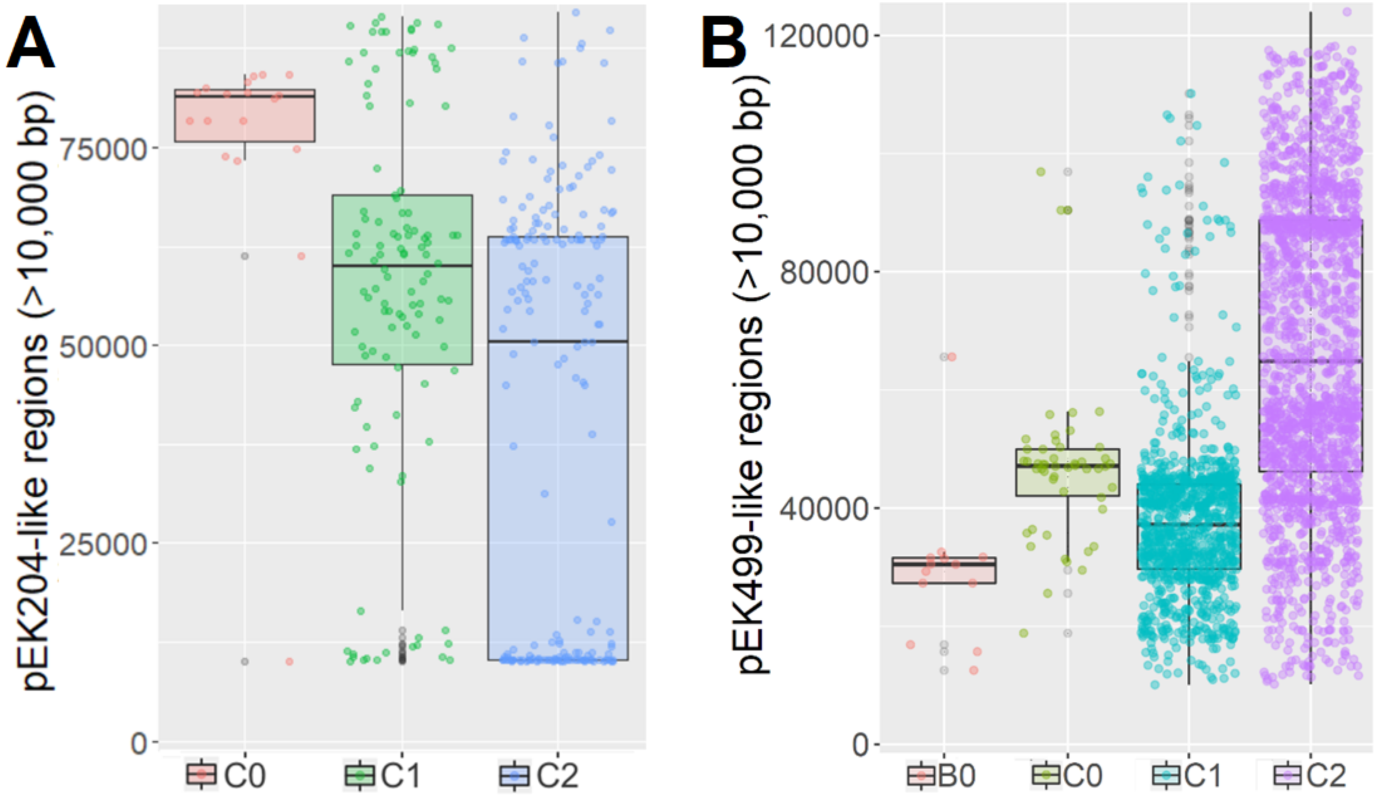


**Figure S7.** The distributions of pEK204-like (A, left) and pEK499-like (B, right) regions in ST131 Clade B and C genome assemblies. Only assemblies with regions spanning >10 Kb in total are shown for matches spanning >300 bp. For pEK204 (93,732 bp), this corresponded to 18 out of 51 for C0 (red), 111 out of 1,119 for C1 (green) and 178 out of 2,051 for C2 (light blue), suggesting initial independent integrations of pEK204-like plasmids across the subclades coupled with subsequent rearrangements. For pEK499 (117,536 bp), this corresponded to 13 out of 14 for B0 (orange), 50 out of 51 for C0 (olive), 1,047 out of 1,119 for C1 (turquoise) and 1,926 out of 2,051 for C2 (mauve), suggesting long-term retention and rearrangement of pEK499-like DNA regions in all four subclades. Note the y-axis length ranges differ.

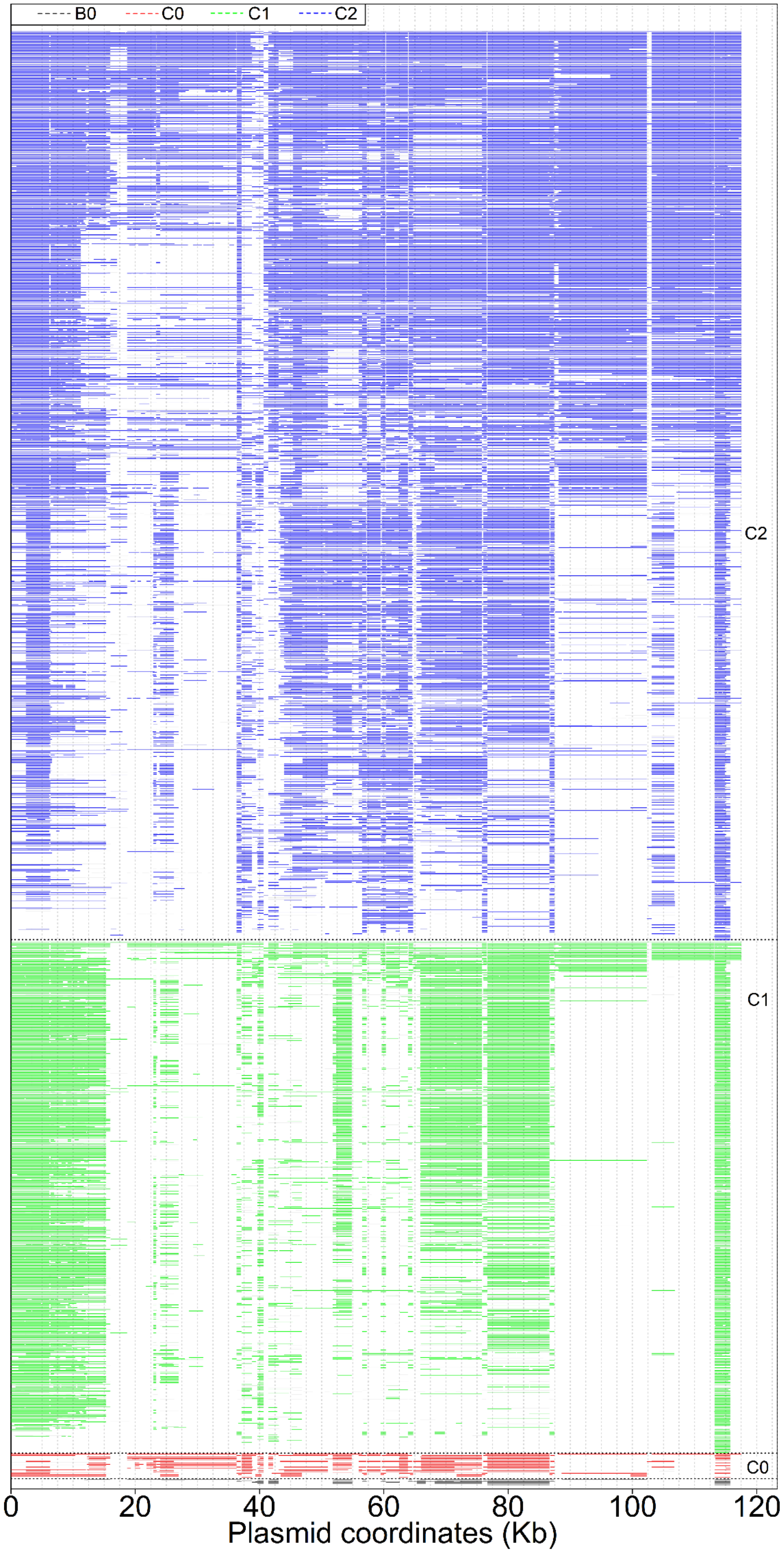


**Figure S8**. The distributions of pEK499-like regions in ST131 Clade B and C genome assemblies. The regions of similarity spanned (from the bottom) B0 (n=14 in black), C0 (n=59 in red), C1 (n=1,047 in green) and C2 (n=1,926 in blue). Similarity across pEK499 (117,536 bp) was based on BLAST alignments. Genes encoding $\mathit{bla}_{\mathrm{TEM}}$, $\mathit{bla}_{\mathrm{OXA\text{-}1}}$ and $\mathit{bla}_{\mathrm{CTX\text{-}M\text{-}15}}$ are at 40, 58 and 63 Kb (respectively).

| Library | Clade | Number of reads | Read length | | |
|---|---|---|---|---|---|
| | | | Media | Mean | SD |
| 8289_1#34_1 | C2 | 2,145,896 | 101 | 99.3 | 6.5 |
| 8289_1#34_2 | | | 100 | 97.7 | 8.0 |
| 8289_1#3_1 | C2 | 1,824,030 | 101 | 99.4 | 6.5 |
| 8289_1#3_2 | | | 100 | 97.6 | 8.2 |
| 8289_1#35_1 | C1 | 2,297,624 | 101 | 99.4 | 6.5 |
| 8289_1#35_2 | | | 100 | 97.7 | 8.0 |
| 8289_1#91_1 | C1 | 2,227,251 | 101 | 99.3 | 6.6 |
| 8289_1#91_2 | | | 100 | 97.7 | 8.1 |
| 8289_1#24_1 | C2 | 2,014,250 | 101 | 99.3 | 6.5 |
| 8289_1#24_2 | | | 100 | 97.7 | 8.1 |
| 8289_1#27_1 | C2 | 5,756,848 | 101 | 99.4 | 6.5 |
| 8289_1#27_2 | | | 100 | 97.7 | 8.1 |
| 8289_1#60_1 | C2 | 1,800,070 | 101 | 99.3 | 6.5 |
| 8289_1#60_2 | | | 100 | 97.6 | 8.1 |
| NCTC13441 | | 2,857,729 | 43 | 42.5 | 1.4 |
| SE15 | | 418,045 | 218 | 192.2 | 67.3 |
| EC958 | | 1,514 | 1,486 | 1,401.6 | 549.2 |

**Table S7**. Read library summary statistics for each main ST131 sample and the three main reference genomes. The read distributions differed for NCTC13441, SE15, and EC958 because they were generated using long read approaches. For GROOT, paired-end read library files were mapped individually. SD stands for standard deviation.

**References not in main text**


Bocher et al. 2009. *Staphylococcus lugdunensis*, a Common Cause of Skin and Soft Tissue Infections in the Community. Journal of Clinical Microbiology, 47(4):946-950.

Kennedy et al. 2010. Complete Nucleotide Sequence Analysis of Plasmids in Strains of *Staphylococcus aureus* Clone USA300 Reveals a High Level of Identity among Isolates with Closely Related Core Genome Sequences. Journal of Clinical Microbiology, 48(12):4504-4511.

Liang et al. 2012. Structural insights into the broadened substrate profile of the extended-spectrum beta-lactamase OXY-1-1 from *Klebsiella oxytoca*. Acta Crystallographica Section D-Biological Crystallography, 68:1460-1467.

Nikaido 2011. Structure and mechanism of rnd-type multidrug efflux pumps. Advances in Enzymology and Related Areas of Molecular Biology, 77:1-60.

Ruiz C, Levy SB. 2010. Many chromosomal genes modulate MarA-mediated multidrug resistance in *Escherichia coli*. Antimicrobial Agents and Chemotherapy, 54(5):2125-2134.

Shaikh S, Fatima J, Shakil S, Rizvi SMD, Kamal MA. 2015. Antibiotic resistance and extended spectrum beta-lactamases: Types, epidemiology and treatment. Saudi Journal of Biological Sciences, 22(1):90-101.

Stoesser N, et al. 2014. Genome sequencing of an extended series of NDM-producing *Klebsiella pneumoniae* isolates from neonatal infections in a Nepali hospital characterizes the extent of community- versus hospital-associated transmission in an endemic setting. Antimicrobial Agents and Chemotherapy, 58(12):7347-7357.

Tse et al. 2010. Complete Genome Sequence of *Staphylococcus lugdunensis* Strain HKU09-01. Journal of Bacteriology, 192(5):1471-1472.
